# Benchmarking reveals superiority of deep learning variant callers on bacterial nanopore sequence data

**DOI:** 10.1101/2024.03.15.585313

**Authors:** Michael B. Hall, Ryan R. Wick, Louise M. Judd, An N. T. Nguyen, Eike J. Steinig, Ouli Xie, Mark R. Davies, Torsten Seemann, Timothy P. Stinear, Lachlan J. M. Coin

**Affiliations:** Department of Microbiology and Immunology, The University of Melbourne, at the Peter Doherty Institute for Infection and Immunity, Melbourne, Australia; Centre for Pathogen Genomics, The University of Melbourne, Melbourne, Victoria, Australia; Department of Infectious Diseases, The University of Melbourne, at the Peter Doherty Institute for Infection and Immunity, Melbourne, Australia; Monash Infectious Diseases, Monash Health, Melbourne, Australia

## Abstract

Variant calling is fundamental in bacterial genomics, underpinning the identification of disease transmission clusters, the construction of phylogenetic trees, and antimicrobial resistance prediction. This study presents a comprehensive benchmarking of SNP and indel variant calling accuracy across 14 diverse bacterial species using Oxford Nanopore Technologies (ONT) and Illumina sequencing. We generate gold standard reference genomes and project variations from closely-related strains onto them, creating biologically realistic distributions of SNPs and indels.

Our results demonstrate that ONT variant calls from deep learning-based tools delivered higher SNP and indel accuracy than traditional methods and Illumina, with Clair3 providing the most accurate results overall. We investigate the causes of missed and false calls, highlighting the limitations inherent in short reads and discover that ONT’s traditional limitations with homopolymer-induced indel errors are absent with high-accuracy basecalling models and deep learning-based variant calls. Furthermore, our findings on the impact of read depth on variant calling offer valuable insights for sequencing projects with limited resources, showing that 10x depth is sufficient to achieve variant calls that match or exceed Illumina.

In conclusion, our research highlights the superior accuracy of deep learning tools in SNP and indel detection with ONT sequencing, challenging the primacy of short-read sequencing. The reduction of systematic errors and the ability to attain high accuracy at lower read depths enhance the viability of ONT for widespread use in clinical and public health bacterial genomics.

## Introduction

Variant calling is a cornerstone of bacterial genomics as well as one of the major applications of next generation sequencing. Its downstream applications include identification of disease transmission clusters, prediction of antimicrobial resistance, and phylogenetic tree construction and subsequent evolutionary analyses, to name a few [1–4]. Variant calling is used extensively in public health laboratories to inform decisions on managing bacterial outbreaks [5] and in molecular diagnostic laboratories as the basis for clinical decisions on how to best treat patients with disease [6].Over the last 15 years, short-read sequencing technologies, such as Illumina, have been the mainstay of variant calling in bacterial genomes, largely due to their relatively high level of base-calling accuracy. However, nanopore sequencing on devices from Oxford Nanopore Technologies (ONT) have emerged as an alternative technology. One of the major advantages of ONT sequencing from an infectious diseases public health perspective is the ability to generate sequencing data in near real-time, as well as the portability of the devices, which has enabled researchers to sequence in remote regions, closer to the source of the disease outbreak [7, 8]. Limitations in ONT basecalling accuracy have historically limited its widespread adoption for bacterial genome variant calling [9]. ONT have recently released a new R10.4.1 pore, along with a new basecaller[10] with three different accuracy modes (fast, high-accuracy [hac] and super-accuracy [sup]). The basecaller also has the ability to identify a subset of paired reads for which both strands have been sequenced (duplex), leading to impressive gains in basecalling accuracy [11–13].

A number of variant callers have been developed for ONT sequencing [14–16]. However, to date, benchmarking studies have focused on human genome variant calling, and have mostly used the older pores, which do not have the ability to identify duplex reads [17–19]. In addition, modern deep learning-based variant callers use models trained on human DNA sequence only, leaving an open question of their generalisability to bacteria [15, 16, 20]. Human genomes have a very different distribution of *k*-mers (segments of DNA sequence of length *k*) and patterns of DNA modification, and as such, results from human studies may not directly carry over into bacterial genomics. Moreover there is substantial *k*-mer and DNA modification variation within bacteria, mandating a broad multi-species approach for evaluation [21]. Existing benchmarks for bacterial genomes, while immensely beneficial and thorough, only assess short-read Illumina data [22, 23].

In this study, we conduct a benchmark of SNP and indel variant calling using ONT and Illumina sequencing across a comprehensive spectrum of 14 Gram-positive and Gram-negative bacterial species. We used the same DNA extractions for both Illumina and ONT sequencing to ensure our results are not biased by acquisition of new mutations during culture. We develop a novel strategy for generating benchmark variant truthsets in which we project variations from different strains onto our gold standard reference genomes in order to create biologically realistic distribution of SNPs and indels. We assess both deep learning-based and traditional variant calling methods and investigate the sources of errors and the impact of read depth on variant accuracy.

## Results

### Genome and variant truthset

Ground truth reference assemblies were generated for each sample using ONT and Illumina reads (see Genome assembly).

Creating a variant truthset for benchmarking is challenging [24, 25]. Calling variants against a sample’s own reference yields no variants, so we generated a mutated reference. Instead of random mutations, we used a pseudo-real approach, applying real variants from a donor genome to the sample’s reference [25, 26]. This approach has the advantage of a simulation, in that we can be certain of the truthset of variants, but with the added benefit of the variants being real differences between two genomes.

For each sample, we selected a donor genome with average nucleotide identity (ANI; a measure of similarity between two genomes) closest to 99.5% (see Truthset and reference generation). We identified all variants between the sample and donor using minimap2 [27] and mummer [28], intersected the variant sets, and removed overlaps and indels longer than 50bp. This variant truthset was then applied to the sample’s reference to create a mutated reference, ensuring no complications from large structural differences. While incorporating structural variation would be an interesting and useful addition to the current work, we chose to focus here on small (<50bp) variants.

***Table 1*** summarises the samples used, the number of variants, and the ANI between each sample and its donor. We analysed 14 samples from different species, spanning a wide range of GC content (30–66%). Despite the variation in SNP counts (2102–57887), the number of indels was consistent across samples (see Suppl. Table S2 for details).

**Table 1.** Summary of the ANI and number of variants found between each sample and its donor genome.

| Sample | Species | ANI (%) | GC (%) | SNPs | Insertions | Deletions | Total variants |
| --- | --- | --- | --- | --- | --- | --- | --- |
| ATCC_33560_202309 | *Campylobacter jejuni* | 99.50 | 30.22 | 6369 | 117 | 106 | 6592 |
| ATCC_35221_202309 | *Campylobacter lari* | 98.64 | 29.81 | 16541 | 57 | 67 | 16665 |
| ATCC_25922_202309 | *Escherichia coli* | 99.50 | 50.42 | 4531 | 119 | 242 | 4892 |
| KPC2_202310 | *Klebsiella pneumoniae* | 99.50 | 57.15 | 15877 | 90 | 78 | 16045 |
| AJ292_202310 | *Klebsiella variicola* | 99.50 | 57.62 | 22850 | 95 | 98 | 23043 |
| ATCC_19119_202309 | *Listeria ivanovii* | 99.46 | 37.13 | 8451 | 187 | 259 | 8897 |
| ATCC_BAA-679_202309 | *Listeria monocytogenes* | 99.50 | 37.98 | 9090 | 66 | 78 | 9234 |
| ATCC_35897_202309 | *Listeria welshimeri* | 99.03 | 36.35 | 16953 | 130 | 133 | 17216 |
| AMtb_1_202402 | *Mycobacterium tuberculosis* | 99.73 | 65.62 | 2102 | 95 | 84 | 2281 |
| ATCC_10708_202309 | *Salmonella enterica* | 99.36 | 52.20 | 18784 | 210 | 189 | 19183 |
| BPH2947_202310 | *Staphylococcus aureus* | 99.48 | 32.80 | 7894 | 95 | 63 | 8052 |
| MMC234_202311 | *Streptococcus dysgalactiae* | 99.16 | 39.49 | 10474 | 82 | 100 | 10656 |
| RDH275_202311 | *Streptococcus pyogenes* | 99.50 | 38.32 | 5361 | 60 | 68 | 5489 |
| ATCC_17802_202309 | *Vibrio parahaemolyticus* | 98.75 | 45.32 | 57887 | 280 | 304 | 58471 |

### Data quality

We analysed ONT data basecalled with three different accuracy models – fast, high accuracy (hac), and super-accuracy (sup) – along with different read types – simplex and duplex (see Basecalling and quality control). Duplex reads are those in which both DNA strands from a single molecule are sequenced back-to-back and basecalled together, whereas simplex reads are basecalled only using a single DNA strand. The median, unfiltered read identities, calculated by aligning reads to their respective assembly, are shown in ***Figure 1***. Duplex reads basecalled with the sup model had the highest median read identity of 99.93% (Q32). The Qscore is the logarithmic transformation of the read identity, *Q* = −10 log_10_ *P*, where *P* is the read identity. This was followed by duplex hac (99.79% [Q27]), simplex sup (99.26% [Q21]), simplex hac (98.31% [Q18]), and simplex fast (94.09% [Q12]). Full summary statistics of the reads can be found in Supplementary Table S1.

**Figure 1.**
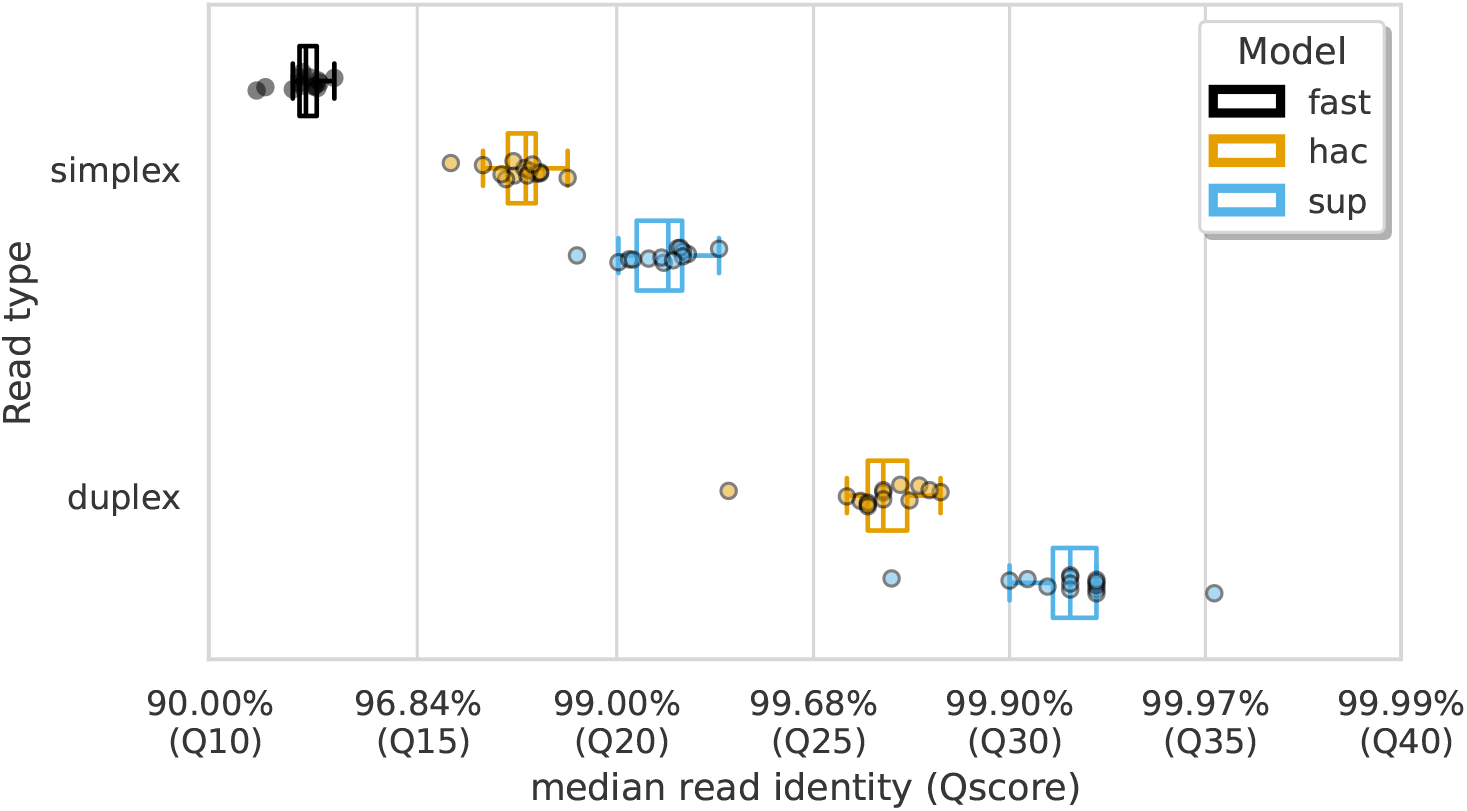
Median alignment-based read identity (x-axis) for each sample (points) stratified by basecalling model (colours) and read type (y-axis). The Qscore is the logarithmic transformation of the read identity, *Q* = −10 log_10_ *P*, where *P* is the read identity.

### Which method is the best?

For this study, we benchmarked the performance of seven variant callers on ONT sequencing data: BCFtools (v1.19 [29]), Clair3 (v1.0.5 [15]), DeepVariant (v1.6.0 [20]), FreeBayes (v1.3.7 [30]), Longshot (v0.4.5 [14]), Medaka(v1.11.3 [31]), and NanoCaller (v3.4.1 [16]). In addition, we called variants from each sample’s Illumina data using Snippy [32] to act as a performance comparison.

Alignments of ONT reads to each sample’s mutated reference (see Genome and variant truth-set) were generated with minimap2 and provided to each variant caller (except Medaka, which takes reads directly). Variant calls were assessed against the truthset using vcfdist (v2.3.3 [33]), classifying each variant as true positive (TP), false positive (FP), or false negative (FN). Precision, recall, and the F1 score were calculated for SNPs and indels at each VCF quality score increment. ***Figure 2*** displays the highest F1 scores for each variant caller across samples, basecalling models, read types, and variant types.

**Figure 2.**
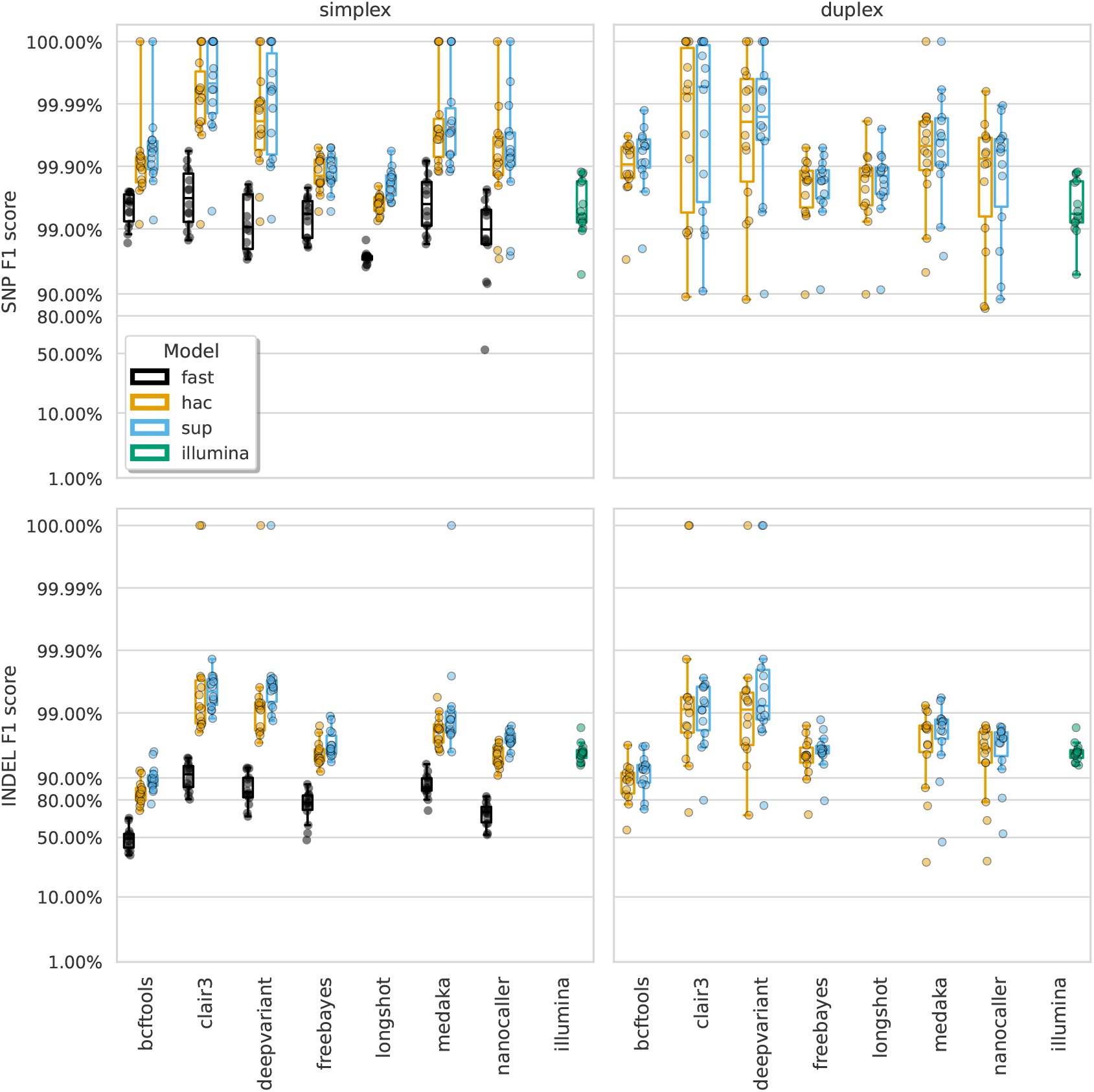
The highest F1 score for each sample (points), stratified by basecalling model (colours), variant type (rows), and read type (columns). Illumina results (green) are included as a reference and do no have different basecalling models or read types. Note, longshot does not provide indel calls.

The F1 score is the harmonic mean of precision and recall and acts as a good metric for overall evaluation. From ***Figure 2*** we see that Clair3 and DeepVariant produce the highest F1 scores for both SNPs and indels with both read types. Unsurprisingly, the sup basecalling model leads to the highest F1 scores across all variant callers, though hac is not much lower. SNP F1 scores of 99.99% are obtained from Clair3 and DeepVariant on sup-basecalled data. For indel calls, Clair3 achieves F1 scores of 99.53% and 99.20% for sup simplex and duplex, respectively, while DeepVariant scores 99.61% and 99.22%. The higher depth of the simplex reads likely explains why the best duplex indel F1 scores are slightly lower than simplex (see How much read depth is enough?). The precision and recall values at the highest F1 score can be seen in Supplementary Figures S3 and S4 (see Suppl. Table S3 for a summary and S4 for full details) as well as results broken down by species for Clair3 with the sup model in Suppl. Figures S5–7. Reads basecalled with the fast model are an order of magnitude worse than the hac and sup models.

***Figure 3*** shows the precision-recall curves for the sup basecalling model (see Suppl. Figures S8 and S9 for the hac and fast model curves, respectively) for each variant and read type – aggregated across samples to produce a single curve for each variant caller. Due to the right-angle-like shape of the Clair3 and DeepVariant curves, filtering based on low-value variant quality improves precision considerably for variant calls, without losing much recall. A similar pattern holds true for BCFtools SNP calls. The best Clair3 and DeepVariant F1 scores are obtained with no quality filtering on sup data, except for indels from duplex data where a quality filter of 4 provides the best F1. See Suppl. Table S5 for the full details.

**Figure 3.**
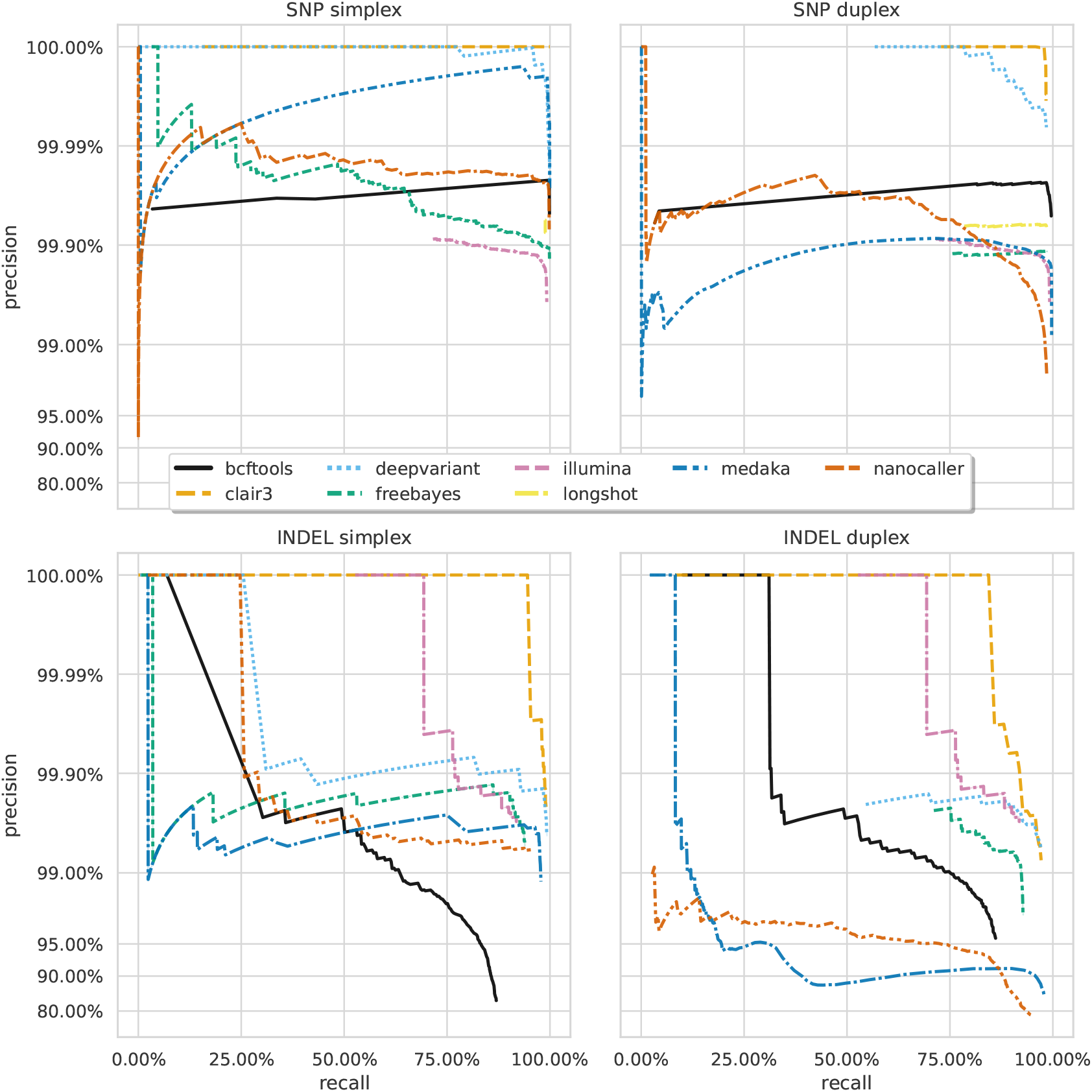
Precision and recall curves for each variant caller (colours and line styles) on sequencing data basecalled with the sup model, stratified by variant type (rows) and read type (columns) and aggregated across samples. The curves are generated by using increasing variant quality score thresholds to filter variants and calculating precision and recall at each threshold. The lowest threshold is the lower right part of the curve, moving to the highest at the top left. Note, Longshot does not provide indel calls.

A striking feature of ***Figure 2*** and ***Figure 3*** is the comparison of deep learning-based variant callers (Clair3, DeepVariant, Medaka, and NanoCaller) to Illumina. For all variant and read types with hac or sup data, these deep learning methods match or surpass Illumina, with median best SNP and indel F1 scores of 99.45% and 95.76% for Illumina. Clair3 and DeepVariant, in particular, perform an order of magnitude better. Traditional variant callers (Longshot, BCFtools, and FreeBayes) match or slightly exceed Illumina for SNP calls with hac and sup data. FreeBayes matches Illumina for indel calls, but BCFtools shows reduced indel accuracy across all models and read types. Fast model ONT data has a lower F1 score than Illumina, only achieving parity in the best case for SNPs.

### Understanding missed and false calls

Conventional wisdom may leave readers surprised at finding that ONT data can provide better variant calls than Illumina. In order to convince ourselves (and others) of these results, we investigate the main causes for this difference.

Given the ONT read-level accuracy now exceeding Q20 (***Figure 1***; simplex sup), read length remains the primary difference between the two technologies. Suppl. Figure S4 shows that Illumina’s lower F1 score is mainly due to recall rather than precision (Suppl. Figure S3). We hypothesised that Illumina errors are related to alignment difficulties in repetitive or variant-dense regions due to its shorter reads.

***Figure 4*** shows that variant density and repetitive regions account for many false negatives, lowering recall. We define variant density as the number of variants (missed or called) in a 100bp window around each call. ***Figure 4***a reveals a bimodal distribution of variant density for Illumina FNs, with a second peak at 20 variants per 100bp, unlike the distribution for TP and FP calls. In contrast, Clair3, a top-performing ONT variant caller, shows no bimodal distribution and few missed or false calls at this density (***Figure 4***b). Illumina reads struggle to align in variant-dense regions, whereas ONT reads can (Suppl. Figure S10), as 20 variants per 100bp represent a larger portion of an Illumina read than an ONT read.

**Figure 4.**
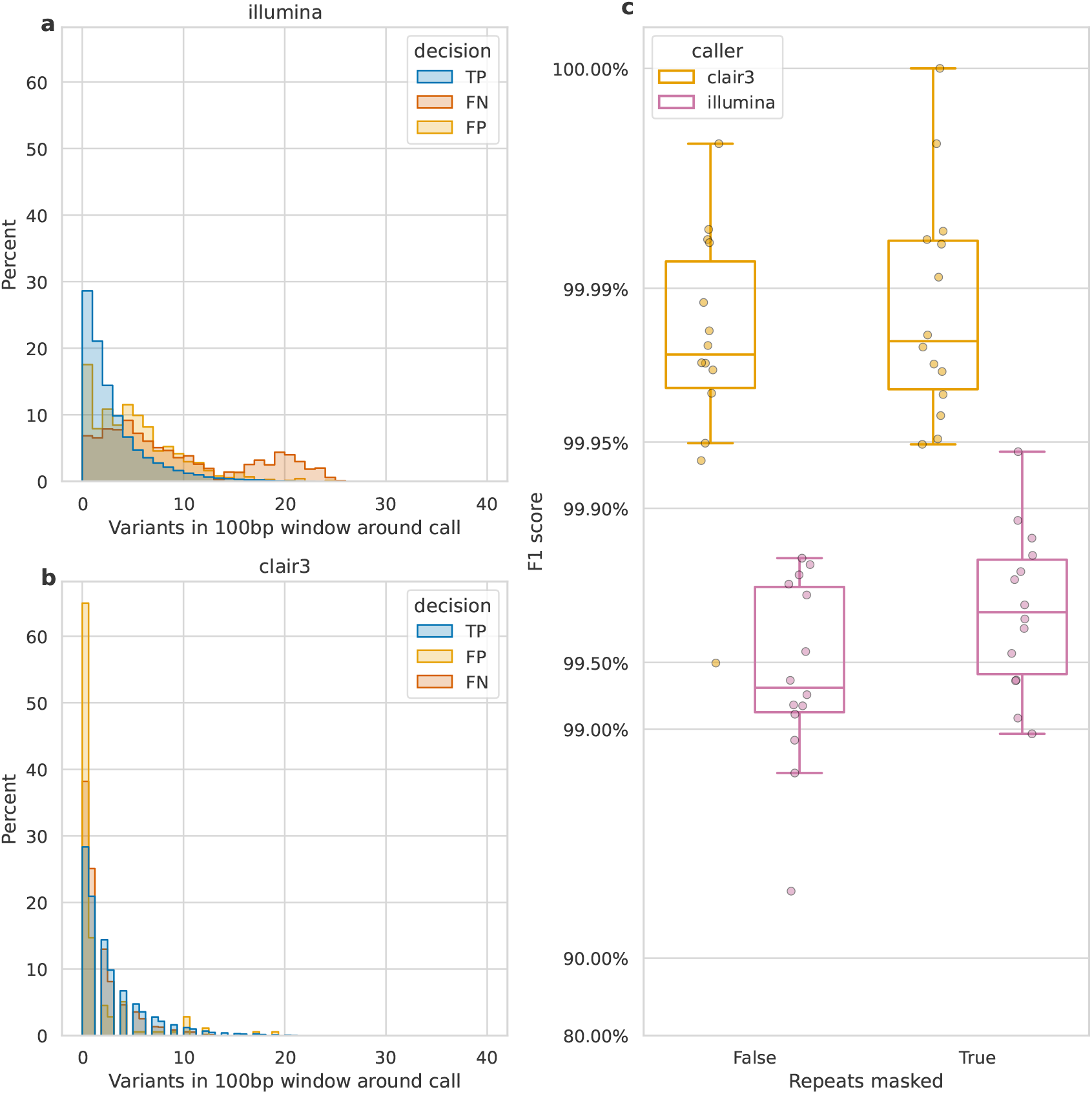
Impact of variant density and repetitive regions on Illumina variant calling. Variant density is the number of (true or false) variants in a 100bp window centred on a call. **a** and **b**) the distribution of variant densities for true positive (TP), false positive (FP) and false negative (FN) calls. The y-axis, percent, indicates the percent of all calls of that decision that fall within the density bin on the x-axis. Illumina calls, aggregated across all samples are shown in **a**, while **b** shows Clair3 calls from simplex sup-basecalled reads at 100x depth. **c**) impact of repetitive regions on the F1 score (y-axis) for Clair3 (100x simplex sup) and Illumina. The x-axis indicates whether variants that fall within repetitive regions are excluded from the calculation of the F1 score. Points indicate the F1 score for a single sample.

We also assessed the change in F1 score when masking repetitive regions of the genome (see Identifying repetitive regions). Due to their shorter length, Illumina reads struggle more with alignment in these regions compared to ONT reads [34]. Suppl. Figure S11 highlights missed variants and alignment gaps in Illumina data. This is further quantified by the increase in Illumina’s F1 score when repetitive regions are masked (***Figure 4***c), rising from 99.3% to 99.7%. In contrast, Clair3 100x simplex sup data shows only a 0.003% increase.

In terms of ONT missed calls, a variant dense repetitive region in the *E. coli* sample ATCC_25922 was the cause of the simplex sup SNP outlier from ***Figure 2*** (see Suppl. Section S2). In addition, the duplex sup SNP outlier was caused by very low read depth for sample KPC2_202310 (*K. pneumoniae*; Suppl. Section S2).

Indels have traditionally been a systematic weakness for ONT sequencing data; primarily driven by variability in the length of homopolymeric regions as determined by basecallers [9]. Having seen the drastic improvements in read accuracy in ***Figure 1***, we sought to determine whether false positive indel calls are still a byproduct of homopolymer-driven errors.

When analysing Clair3, the best-performing ONT caller, we found that reads basecalled with the fast model often miscalculate homopolymer lengths by 1 or 2bp (***Figure 5***), though there is an equal number of non-homopolymeric false indel calls. In contrast, the sup model significantly reduced false indel calls, matching Illumina’s error profile. Of the eight false indel calls by Clair3 on sup data, five were homopolymers and three occurred within one or two bases of another insertion with a similar sequence. The hac model improved over the fast model but still produced notable false indel calls, mainly miscalculating homopolymers by 1bp. DeepVariant showed a similar error profile to Clair3 (Suppl. Figure S13), with 8/11 false indels being homopolymers. FreeBayes (Suppl. Figure S14), Medaka (Suppl. Figure S15) and NanoCaller (Suppl. Figure S16) performed similarly, while BCFtools (Suppl. Figure S12) exhibited a persistent bias for homopolymeric indel errors, even with sup model reads. This indicates that while the sup basecaller reduces bias, deep learning methods like Clair3 and DeepVariant further mitigate it by training models to account for these systematic issues. An honourable mention goes to FreeBayes, a traditional variant caller that handles errors without inherent bias.

**Figure 5.**
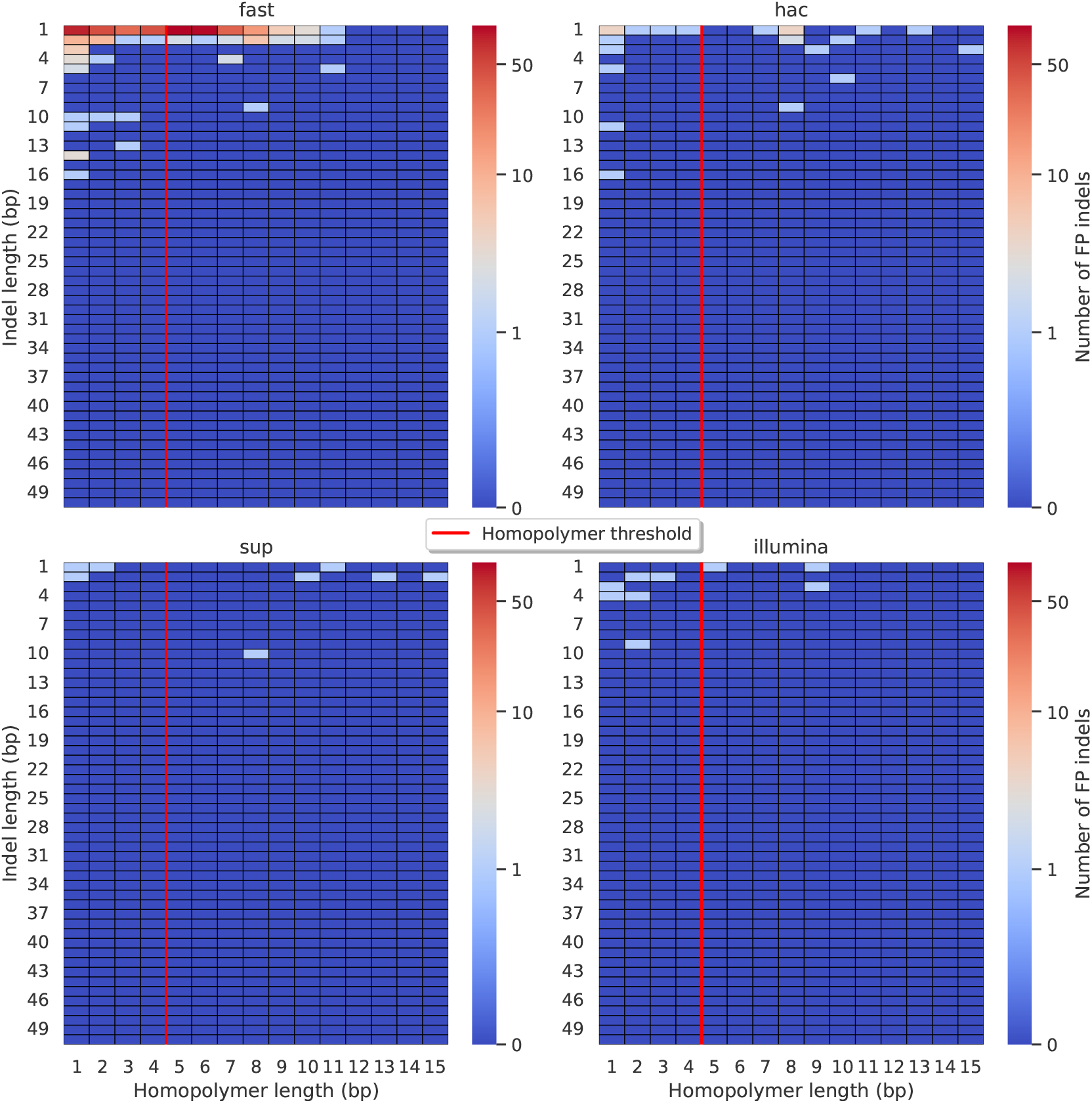
Relationship between indel length (y-axis) and homopolymer length (x-axis) for false positive (FP) indel calls for Clair3 100x simplex fast (top left), hac (top right), and sup (lower left) calls. Illumina is shown in the lower right for reference. The vertical red line indicates the threshold above which we deem a run of the same nucleotide to be a ‘true’ homopolymer. Indel length is the number of bases inserted/deleted for an indel, whereas the homopolymer length indicates how long the tract of the same nucleotide is after the indel. The colour of a cell indicates how many FP indels of that indel-homopolymer length combination.

Lastly, we did not see any systematic indel bias in the context of missed calls (Suppl. Figures S17–S22), especially when compared to Illumina indel error profiles.

### How much read depth is enough?

Having established the accuracy of variant calls from ‘full depth’ ONT datasets (100x), we investigated the required ONT read depth to achieve desired precision or recall, which varies by use case and resource availability. This is particularly relevant for ONT, where sequencing can be stopped in real-time once ‘sufficient’ data is obtained.

We subsampled each ONT read set with rasusa (v0.8.0 [35]) to average depths of 5, 10, 25, 50, and 100x and called variants with these reduced sets. Due to limited duplex depth, 50x was the maximum used for duplex reads, while 100x was used for simplex reads.

***Figure 6*** and ***Figure 7*** show F1 score, precision, and recall as functions of read depth for SNPs and indels. Precision and recall decrease as read depth is reduced, notably below 25x. Remarkably, Clair3 or DeepVariant on 10x ONT sup simplex data provides F1 scores consistent with, or better than, full-depth Illumina for both SNPs and indels (see Table S1 for Illumina read depths). The same is true for duplex hac or sup reads.

**Figure 6.**
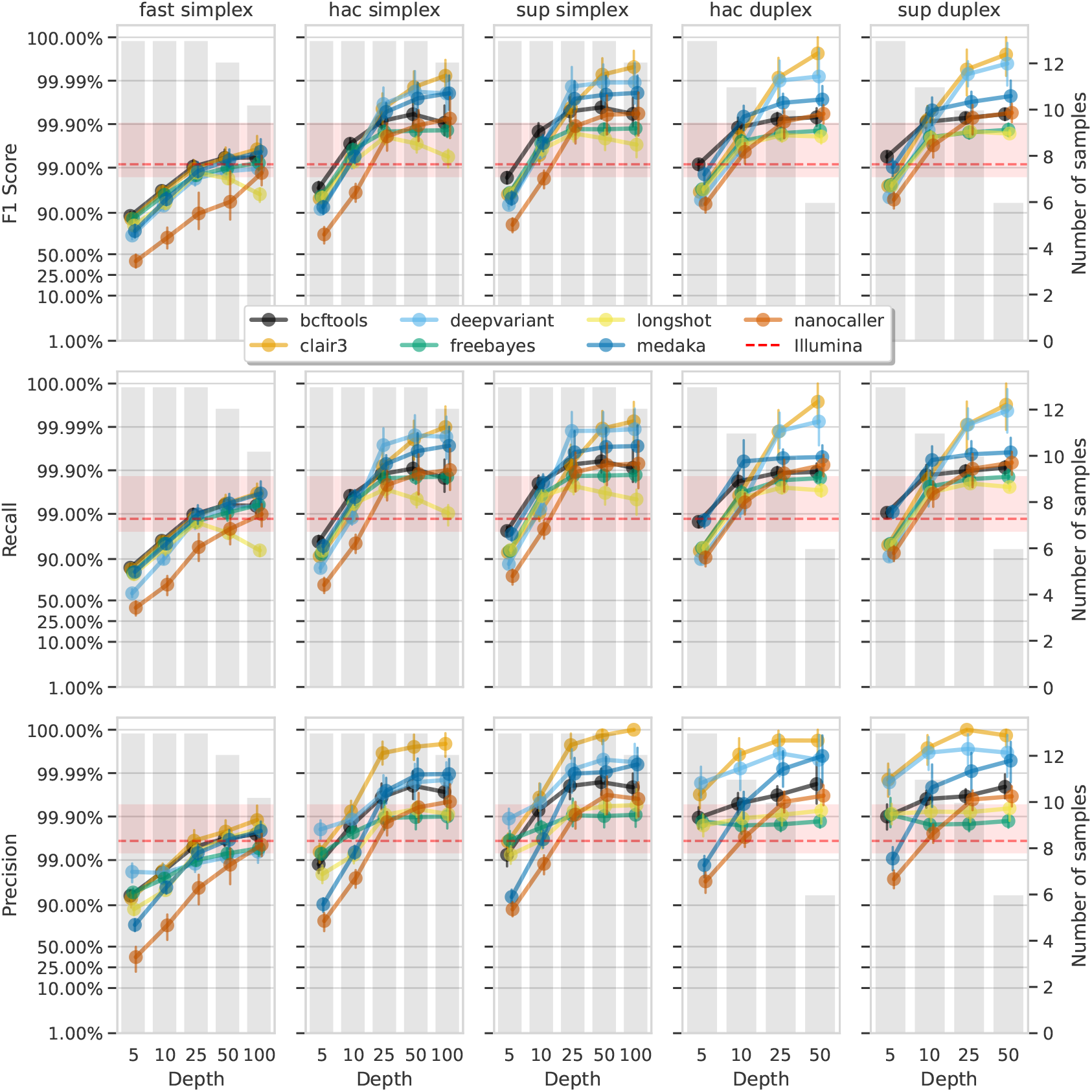
Effect of read depth (x-axis) on the highest SNP F1 score, and precision and recall at that F1 score (y-axis), for each variant caller (colours). Each column is a basecall model and read type combination. The grey bars indicate the number of samples with at least that much read depth in the full read set. Samples with less than that depth were not used to calculate that depth’s metrics. Bars on each point at each depth depict the 95% confidence interval. The horizontal red dashed line is the full-depth Illumina value for that metric, with the red bands indicating the 95% confidence interval.

**Figure 7.**
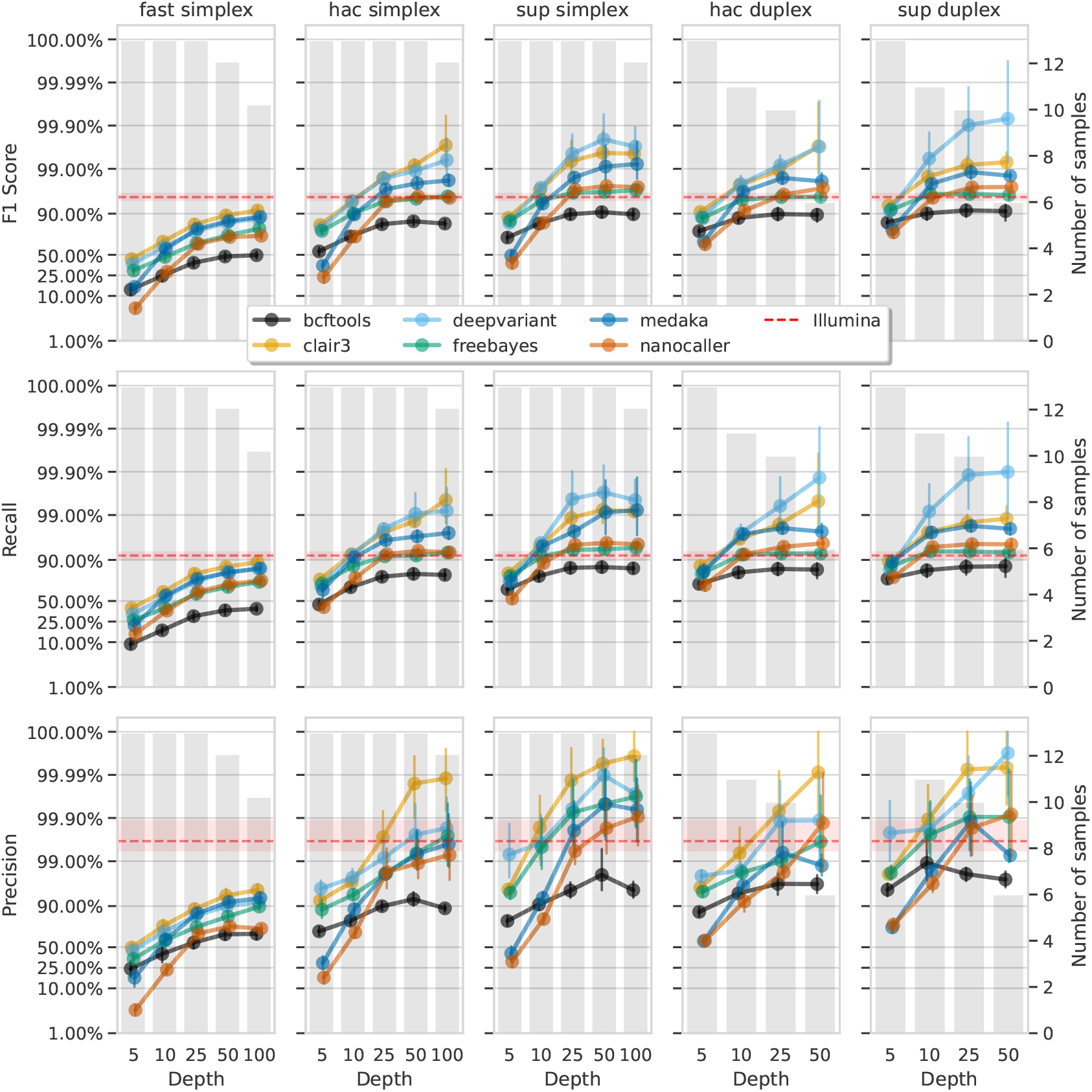
Effect of read depth (x-axis) on the highest indel F1 score, and precision and recall at that F1 score (y-axis), for each variant caller (colours). Each column is a basecall model and read type combination. The grey bars indicate the number of samples with at least that much read depth in the full read set. Samples with less than that depth were not used to calculate that depth’s metrics. Bars on each point at each depth depict the 95% confidence interval. The horizontal red dashed line is the full-depth Illumina value for that metric, with the red bands indicating the 95% confidence interval.

With 5x of ONT read depth the F1 score is lower than Illumina for almost all variant caller and basecalling models. However, BCFtools surprisingly produces SNP F1 scores on par with Illumina on duplex sup reads. Despite the inferior F1 scores across the board at 5x, SNP precision remains above Illumina with duplex reads for all methods except NanoCaller, and calls from Clair3 and DeepVariant simplex sup data.

### What computational resources do I need?

The final consideration for variant calling is the required computational resources. While this may be trivial for those with high-performance computing (HPC) access, many analyse bacterial genomes on personal computers due to their smaller size compared to eukaryotes. The main resource constraints are memory and runtime, especially for aligning reads to a reference and calling variants. Additionally, if working with raw (pod5) ONT data, basecalling is also a resource-intensive step.

***Figure 8*** shows the runtime (seconds per megabase of sequencing data) and maximum memory usage for read alignment and variant calling (see Suppl. Figure S23 and Table S7 for basecalling GPU runtimes). DeepVariant was the slowest (median 5.7s/Mbp) and most memory-intensive (median 8GB), with a runtime of 38 minutes for a 4Mbp genome at 100x depth. FreeBayes had the largest runtime variation, with a maximum of 597s/Mbp, equating to 2.75 days for the same genome. In contrast, basecalling with a single GPU using the super-accuracy model required a median runtime of 0.77s/Mbp, or just over 5 minutes for a 4Mbp genome at 100x depth. Clair3 had a median memory usage of 1.6GB and a runtime of 0.86s/Mbp (<6 minutes for a 4Mbp 100x genome). Full details are in Suppl. Table S6.

**Figure 8.**
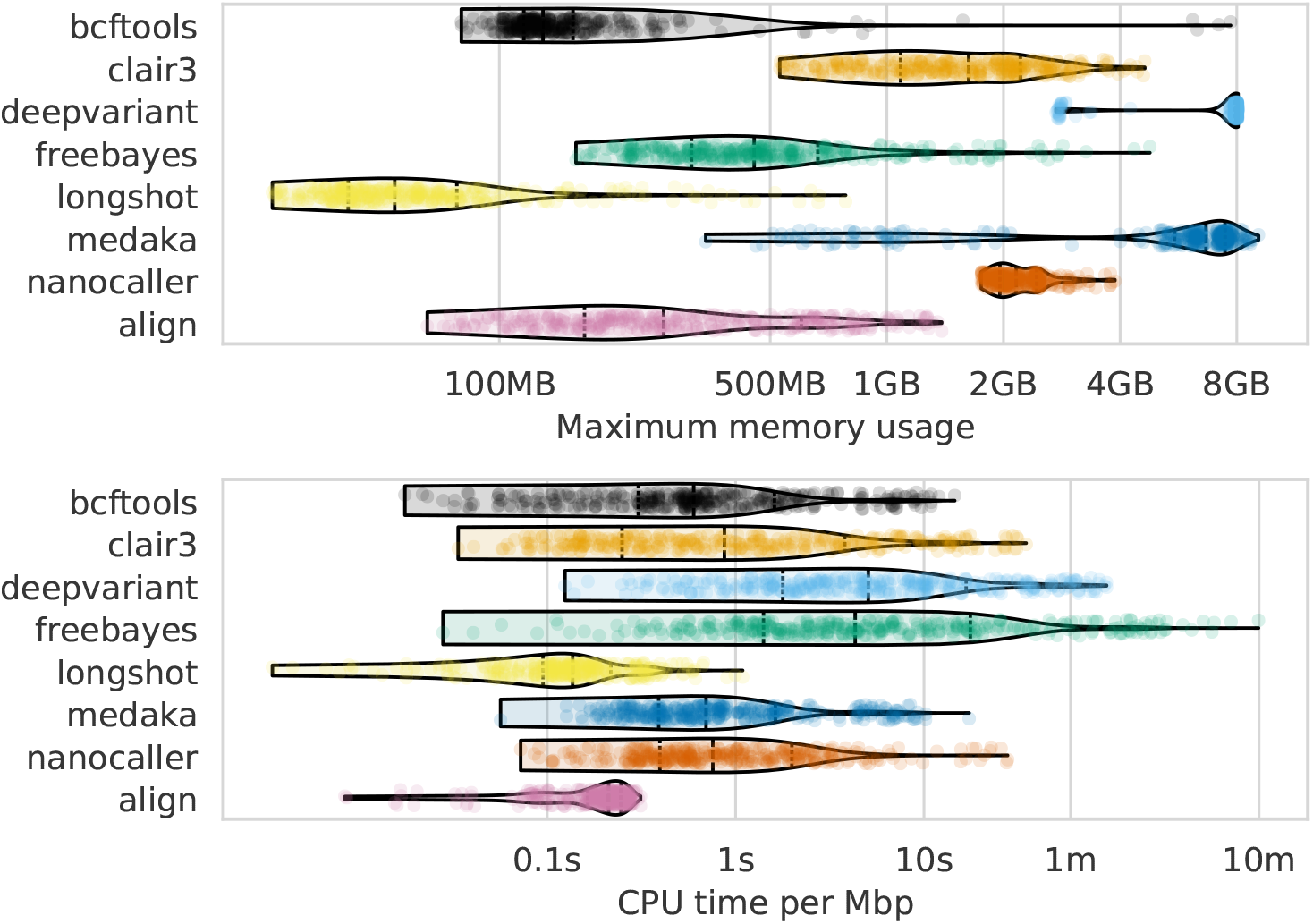
Computational resource usage of alignment and each variant caller (y-axis and colours). The top panel shows the maximum memory usage (x-axis) and the lower panel shows the runtime as a function of the CPU time (seconds) divided by the number of basepairs in the readset (seconds per megabasepairs; x-axis). Each point represents a single run across read depths, basecalling models, read types, and samples for that variant caller (or alignment). s=seconds; m=minutes; MB=megabytes; GB=gigabytes; Mbp=megabasepairs.

## Discussion

In this study, we evaluated the accuracy of bacterial variant calls derived from Oxford Nanopore Technologies (ONT) using both conventional and deep learning-based tools. Our findings show that deep learning approaches, specifically Clair3 and DeepVariant, deliver high accuracy in SNP and indel calls from the latest high-accuracy basecalled ONT data, outperforming Illumina-based methods, with Clair3 achieving median F1 scores of 99.99% for SNPs and 99.53% for indels.

Our dataset comprised deep sequencing of 14 bacterial species using the latest ONT R10.4.1 flowcells, with a 5 kHz sampling rate, and complementary deep Illumina sequencing. Consistent with previous studies [11–13], we observed read accuracies greater than 99.0% (Q20) and 99.9% for simplex and duplex reads, respectively (***Figure 1***).

The high-quality sequencing data enabled the creation of near-perfect reference genomes, crucial for evaluating variant calling accuracy. While not claiming perfection for these genomes, we consider them to be as accurate as current technology allows (or as philosophically possible) [13, 36].

To benchmark variant calling, we utilised a variant truthset generated by applying known differences between closely related genomes to a reference. This pseudo-real method offers a realistic evaluation framework and a reliable truthset for assessing variant calling accuracy [25, 26].

Our comparison of variant calling methods showed that deep learning techniques achieved the highest F1 scores for SNP and indel detection, indicating their potential in genomic analyses and suggesting a shift towards more advanced computational approaches. While the superior performance of these methods has been established for human variant calls [17, 18], our results confirm their effectiveness for bacterial genomes as well.

Our investigation into missed and false variant calls highlights inherent challenges posed by sequencing technology limitations, particularly read length, alignment in complex regions, and indel length in homopolymers. We found that variant density and repetitive regions hinder Illumina variant calling due to short read alignment issues. However, we found recent improvements in ONT read accuracy and deep learning-based variant callers have mitigated homopolymer-induced false positive indel calls, previously a major systematic issue with ONT data [9, 13].

Having established the accuracy and error sources of modern methods, we examined the impact of read depth on variant calling accuracy. Our results show that high accuracy is achievable at reduced read depths of 10x, especially with super-accuracy basecalling models and deep learning algorithms. This is significant for resource-limited projects, as 10x super-accuracy simplex data can match or exceed Illumina accuracy. For optimal clinical and public health applications, we recommend a minimum of 25x depth. Notably, 5x depth with duplex super-accuracy ONT data achieved SNP accuracy comparable to Illumina. Having such confidence in low-depth calls will no doubt be a boon for many clinical and public health applications where sequencing direct-from-sample is desired [2, 37–39].

Lastly, considering computational resource requirements is crucial, especially for those without high-performance computing facilities [7, 8, 40]. Our findings show a wide range of demands among variant calling methods, with the worst-case scenario (FreeBayes) taking over two days. Most methods, however, run in less than 40 minutes, with Clair3 having a median runtime of about 6 minutes. All methods use less than 8GB of memory, making them compatible with most laptops. Basecalling is generally faster than variant calling, assuming GPU access, which is likely considered when acquiring ONT-related equipment.

There are three main limitations to this work. The first is that we only assess small variants and ignored structural variants. Zhou *et al*. benchmarked structural variant calling from ONT data [41], though this focused on human sequencing data. Generating a truthset of structural variants between two genomes is, in itself, a difficult task. However, we believe such an undertaking with a thorough investigation of structural variant calling methods for bacterial genomes would be highly beneficial.

The second limitation is not using a diverse range of ANI values for selecting the variant donor genomes when generating the truthset. Previous work from Bush *et al*. examined different diversity thresholds for selecting reference genomes when calling variants from Illumina data, and found it to be one of the main differentiating factors in accuracy [23]. Our results mirror this to an extent, showing the reduction in Illumina accuracy as the variant density increases, though it would be interesting to determine whether the divergence in reference genomes has an affect on ONT variant calling accuracy. Nevertheless, to maintain our focus on the nuances of variant calling methods, including basecalling models, read types, error types, and the influence of read depth, we decided that introducing another layer of complexity into our benchmark could potentially obscure some of the insights.

The third limitation is that Illumina sequencing was performed on different models: three samples on the NextSeq 500 and the rest on the NextSeq 2000. While differences in error rates exist between Illumina instruments, no specific assessment has been made between these NextSeq models [42]. However, the absolute differences in error rates are minor and unlikely to impact our study significantly. This is particularly relevant since Illumina’s lower F1 score compared to ONT was due to missed calls rather than erroneous ones.

In conclusion, this study comprehensively evaluates bacterial variant calls using Oxford Nanopore Technologies (ONT), highlighting the superior performance of deep learning tools, particularly Clair3 and DeepVariant, in SNP and indel detection. Our extensive dataset and rigorous benchmarking demonstrate significant advancements in sequencing accuracy with the latest ONT technologies.

Improvements in ONT read accuracy and deep learning variant callers have mitigated previous challenges like homopolymer-associated errors. We also found that high accuracy can be achieved at lower read depths, making these methods practical for resource-limited settings. This capability marks a significant step in making advanced genomic analysis more accessible and impactful.

## Methods

### Sequencing

Bacterial isolates were streaked onto agar plates and grown overnight at 37°C. *Mycobacterium tu-berculosis, Streptococcus pyogenes*, and *Streptococcus dysgalatiae* subsp. *equisimilis* were grown in liquid media of 7H9 or TSB with shaking until reaching high cell density (OD ∼ 1; see Suppl. Section S1 for *Streptococcus* sample selection). The cultures were centrifuged at 13000rpm for 10 minutes and cell pellets were collected. Bacteria were lysed with appropriate enzymatic treatment except for *Mycobacterium* and *Streptococcus*, which were lysed by bead beating (PowerBead, 0.5mm glass beads [13116-50] or Lysing Matrix Y [116960050-CF] and Precellys or Tissue lyser [Qiagen]). DNA extraction was performed by sodium acetate precipitation and further Ampure XP bead purification (Beckman Coulter) or either Beckman Coulter GenFind V2 (A41497) or QIAGEN Blood and Tissue DNEasy kit (69506). Illumina library preparation was performed using Illumina DNA prep (20060059) using quarter reagents and Illumina DNA/RNA UD Indexes. Short-read whole-genome sequencing was performed on an Illumina NextSeq 500 for the *M. tuberculosis* (AMtb_1 202402), *S. pyogenes* (RDH275 202311), and *S. dysgalactiae* (MMC234 202311) samples and a NextSeq 2000 for all other samples, with a 150bp PE kit. ONT library preparation was performed using either Rapid Barcoding Kit V14 (SQK-RBK114.96) or Native Barcoding Kit V14 (SQK-NBD114.96). Long-read whole-genome sequencing was performed on a MinION Mk1b or GridION using R10.4.1 MinION flow cells (FLO-MIN114). Supplementary Table S9 contains detailed information about the instru-ment models and multiplexing for each sample.

### Basecalling and quality control

Raw ONT data were basecalled with Dorado (v0.5.0 [10]) using the v4.3.0 models *fast* (dna_r10.4.1 _e8.2_400bps_fast@v4.3.0), *hac* (dna_r10.4.1_e8.2_400bps_hac@v4.3.0), and *sup* (dna_r10.4.1_e8.2 _400bps_sup@v4.3.0). Duplex reads were additionally generated using the duplex subcommand of Dorado with hac and sup models (fast is not compatible with duplex). All runs were basecalled on an Nvidia A100 GPU to ensure consistency. Reads shorter than 1000bp or with a quality score below 10 were removed with SeqKit (v2.6.1 [43]) and each readset was randomly subsampled to a maximum mean read depth of 100x with Rasusa (v0.8.0 [35]).

Illumina reads were preprocessed with fastp (v0.23.4 [44]) to remove adapter sequences, trim low-quality bases from the ends of the reads, and remove duplicate reads and reads shorter than 30bp.

### Genome assembly

Ground truth assemblies were generated for each sample as per Wick *et al*. [36]. Briefly, the unfiltered ONT simplex sup reads were filtered with Filtlong (v0.2.1 [45]) to keep the best 90% (-p 90) and fastp (default settings) was used to process the raw Illumina reads. We performed 24 separate assemblies using the *Extra-thorough assembly* instructions in Trycycler’s (v0.5.4 [46]) documentation. Assemblies were combined into a single consensus assembly with Trycycler and Illumina reads were used to polish that assembly using Polypolish (v0.6.0; default settings [47]) and Pypolca (v0.3.1 [48, 49]) with `--careful`. Manual curation and investigation of all polishing changes was made as per Wick *et al*. [36] (e.g., for very long homopolymers, the correct length was chosen as per Illumina reads support).

### Truthset and reference generation

To generate the variant truthset for each sample, we identified all variants between the sample and a *donor* genome. To select the variant-donor genome for a given sample, we downloaded all RefSeq assemblies for that species (up to a maximum of 10000) using genome_updater (v0.6.3 [50]). ANI was calculated between each downloaded genome and the sample reference using skani (v0.2.1 [51]). We only kept genomes with an ANI, *a*, such that 98.40% ≤ *a* <= 99.80%. In addition, we excluded any genomes with CheckM[52] completeness less than 98% and contamination greater than 5%. We then selected the genome with the ANI closest to 99.50%. Our reasoning for this range exclusion is that genomes with *a* < 99.80% are almost always members of the same sequence type (ST)[53, 54], and we found very little variation between them (data not shown).

We then identified variants between the reference and donor genomes using both minimap2 (v2.26 [27]) and mummer (v4.0.0rc1 [28]). We took the intersection of the variants identified by minimap2 and mummer into a single VCF and used BCFtools (v1.19 [29]) to decompose multi nucleotide polymorphisms (MNPs) into SNPs, left-align and normalise indels, remove duplicate and overlapping variants, and exclude any indel longer than 50bp. The resulting VCF file is our truthset.

Next, we generated a mutated reference genome, which we used as the reference against which variants were called by the different methods we assess. BCFtools’ consensus subcommand was used to apply the truthset of variants to the sample reference, thus producing a mutated reference.

### Alignment and variant calling

ONT reads were aligned to the mutated reference with minimap2 using options `--cs, --MD,` and `-aLx map-ont` and output to a BAM alignment file.

Variant calling was performed from the alignment files with BCFtools (v1.19 [29]), Clair3 (v1.0.5 [15]), DeepVariant (v1.6.0 [20]), FreeBayes (v1.3.7 [30]), Longshot (v0.4.5 [14]), and NanoCaller (v3.4.1 [16]). In addition, variant calling was performed directly from the reads for Medaka (v1.11.3 [31]) as Medaka does its own alignment with minimap2. Individual parameters used for each variant caller can be found in the accompanying GitHub repository [55, 56].

Where a variant caller provided an option to set the expected ploidy, haploid was given. In addition, where a minimum read depth or base quality option was available, a value of 2 and 10, respectively, was used in order to try and make downstream assessment and filtering consistent across callers.

For Clair3, the pretrained models for Dorado model v4.3.0 provided by ONT were used [57]. However, as no fast model is available, we used the hac model with the fast-basecalled reads. The pretrained model option `--model_type ONT_R104` was used with DeepVariant, and the default model was used for NanoCaller. For Medaka, the provided v4.3.0 sup and hac models were used, with the hac model being used for fast data as no fast model is available.

For the Illumina variant calls that act as a benchmark to compare ONT against, we chose Snippy [32] due to being tailored for haploid genomes and being one of the best performing variant callers on Illumina data [23]. Snippy performs alignmen of read with BWA-MEM [58] and calls variants with FreeBayes.

Variant call files (VCFs) are then filtered to remove overlapping variants, make heterzygous calls homozygous for the allele with the most depth, normalise and left-align indels, break MNPs into SNPs and remove indels longer than 50bp, all with BCFtools.

### Variant call assessment

Filtered VCFs were assessed with vcfdist (v2.3.3 [33]) using the truth VCFs and mutated references from Truthset and reference generation. We disabled partial credit with `--credit-threshold 1.0` and set the maximum variant quality threshold (`-mx`) to the maximum in the VCF being assessed.

### Identifying repetitive regions

To identify repetitive regions in the mutated reference, we used the following mummer utilities. `nucmer --maxmatch --nosimplify` to align the reference against itself and retain non-unique alignments. We then passed the output into `show-coords -rTH -I 60` to obtain the coordinates for all alignments with an identity of 60% or greater. Alignments where the start and end coordinates of the alignment do not match are considered as repeats and these are output in the BED format, with intervals being merged with BEDtools [59].

## Code availability

All code to perform the analyses in this work are available on GitHub and archived on Zenodo [55, 56].

## Data availability

The unfiltered fastq, and assembly files generated in this study have been submitted to the NCBI BioProject database under accession numbers PRJNA1087001 and PRJNA1042815. See Suppl. Table S8 for a list of all Assembly, BioSample and Run accessions, as well as DOIs for raw ONT pod5 data. Variant truthsets and associated data are archived on Zenodo [60].

## Acknowledgements

This research was performed in part at Doherty Applied Microbial Genomics, Department of Microbiology and Immunology, The University of Melbourne at the Peter Doherty Institute for Infection and Immunity. This research was supported by the University of Melbourne’s Research Computing Services and the Petascale Campus Initiative. We are grateful to Bart Currie and Tony Korman for providing the *Streptococcus dysgalactiae* (MMC234 202311) and *Streptococcus pyogenes* (RDH275 202311) samples. We thank Zamin Iqbal and Martin Hunt for insightful discussions relating to truthset and reference generation. We would also like to thank Romain Guérillot and Miranda Pitt for invaluable feedback throughout the project.

## Author contributions

**Michael B. Hall:** Conceptualization, Data curation, Formal Analysis, Investigation, Methodology, Project administration, Software, Writing – original draft, Visualization, Writing – review & editing. **Ryan R. Wick:** Conceptualization, Data curation, Formal Analysis, Investigation, Methodology, Software, Writing – original draft, Writing – review & editing. **Louise M. Judd:** Conceptualization, Data curation, Investigation, Methodology, Project administration, Resources, Writing – original draft, Writing – review & editing. **An N. T. Nguyen:** Investigation, Methodology, Resources, Writing – original draft, Writing – review & editing. **Eike J. Steinig:** Conceptualization, Data curation, Formal Analysis, Investigation, Methodology, Software, Writing – review & editing. **Ouli Xie:** Methodology, Resources, Writing – review & editing. **Mark R. Davies:** Methodology, Resources, Supervision, Writing – review & editing. **Torsten Seemann:** Conceptualization, Methodology, Supervision, Writing – review & editing. **Lachlan J. M. Coin:** Conceptualization, Funding acquisition, Methodology, Resources, Supervision, Writing – original draft, Writing – review & editing. **Timothy P. Stinear:** Conceptualization, Funding acquisition, Methodology, Resources, Supervision, Writing – review & editing.

## References

[1] James Stimson et al. ‘Beyond the SNP Threshold: Identifying Outbreak Clusters Using Inferred Transmissions’. In: Molecular Biology and Evolution 36.3 (2019), pp. 587–603. ISSN: 0737-4038. DOI: 10.1093/molbev/msy242.

[2] Dropen Sheka, Nikolay Alabi, and Paul MK Gordon. ‘Oxford nanopore sequencing in clinical microbiology and infection diagnostics’. In: Briefings in Bioinformatics 22.5 (2021), bbaa403. ISSN: 1477-4054. DOI: 10.1093/bib/bbaa403.

[3] Timothy M Walker et al. ‘The 2021 WHO catalogue of Mycobacterium tuberculosis complex mutations associated with drug resistance: a genotypic analysis’. In: The Lancet Microbe (2022). ISSN: 2666-5247. DOI: 10.1016/s2666-5247(21)00301-3.

[4] Frederic Bertels et al. ‘Automated Reconstruction of Whole-Genome Phylogenies from Short-Sequence Reads’. In: Molecular Biology and Evolution 31.5 (2014), pp. 1077–1088. ISSN: 0737-4038. DOI: 10.1093/molbev/msu088.

[5] Claire L Gorrie et al. ‘Key parameters for genomics-based real-time detection and tracking of multidrug-resistant bacteria: a systematic analysis’. In: The Lancet Microbe (2021). ISSN: 2666-5247. DOI: 10.1016/s2666-5247(21)00149-x.

[6] Norelle L. Sherry et al. ‘An ISO-certified genomics workflow for identification and surveillance of antimicrobial resistance’. In: Nature Communications 14.1 (2023), p. 60. ISSN: 2041-1723. DOI: 10.1038/s41467-022-35713-4.

[7] Nuno Rodrigues Faria et al. ‘Mobile real-time surveillance of Zika virus in Brazil’. In: Genome Medicine 8.1 (2016), p. 97. DOI: 10.1186/s13073-016-0356-2.

[8] Thomas Hoenen et al. ‘Nanopore Sequencing as a Rapidly Deployable Ebola Outbreak Tool’. In: Emerging Infectious Diseases 22.2 (2016), pp. 331–334. ISSN: 1080-6040. DOI: 10.3201/eid2202.151796.

[9] Clara Delahaye and Jacques Nicolas. ‘Sequencing DNA with nanopores: Troubles and biases’. In: PLOS ONE 16.10 (2021), e0257521. ISSN: 1932-6203. DOI: 10.1371/journal.pone.0257521.

[10] Oxford Nanopore Technologies. Dorado: Oxford Nanopore’s Basecaller. Version 0.5.0. 2023. URL: https://github.com/nanoporetech/dorado.

[11] Nicholas D. Sanderson et al. ‘Comparison of R9.4.1/Kit10 and R10/Kit12 Oxford Nanopore flowcells and chemistries in bacterial genome reconstruction’. In: Microbial Genomics 9.1 (2023), p. 000910. ISSN: 2057-5858. DOI: 10.1099/mgen.0.000910.

[12] Nicholas D. Sanderson et al. ‘Evaluation of the accuracy of bacterial genome reconstruction with Oxford Nanopore R10.4.1 long-read-only sequencing’. In: bioRxiv (2024). DOI: 10.1101/2024.01.12.575342.

[13] Mantas Sereika et al. ‘Oxford Nanopore R10.4 long-read sequencing enables the generation of near-finished bacterial genomes from pure cultures and metagenomes without short-read or reference polishing’. In: Nature Methods (2022), pp. 1–4. ISSN: 1548-7091. DOI: 10.1038/s41592-022-01539-7.

[14] Peter Edge and Vikas Bansal. ‘Longshot enables accurate variant calling in diploid genomes from single-molecule long read sequencing’. In: Nature Communications 10.1 (2019), p. 4660. ISSN: 2041-1723. DOI: 10.1038/s41467-019-12493-y.

[15] Zhenxian Zheng et al. ‘Symphonizing pileup and full-alignment for deep learning-based long-read variant calling’. In: Nature Computational Science 2.12 (2022), pp. 797–803. ISSN: 2662-8457. DOI: 10.1038/s43588-022-00387-x.

[16] Mian Umair Ahsan et al. ‘NanoCaller for accurate detection of SNPs and indels in dificult-to-map regions from long-read sequencing by haplotype-aware deep neural networks’. In: Genome Biology 22.1 (2021), p. 261. ISSN: 1474-760X. DOI: 10.1186/s13059-021-02472-2.

[17] Nathan D. Olson et al. ‘Variant calling and benchmarking in an era of complete human genome sequences’. In: Nature Reviews Genetics 24.7 (2023), pp. 464–483. ISSN: 1471-0064. DOI: 10.1038/s41576-023-00590-0.

[18] Nathan D. Olson et al. ‘PrecisionFDA Truth Challenge V2: Calling variants from short and long reads in dificult-to-map regions’. In: Cell Genomics 2.5 (2022), p. 100129. ISSN: 2666-979X. DOI: 10.1016/j.xgen.2022.100129.

[19] Surui Pei et al. ‘Benchmarking variant callers in next-generation and third-generation sequencing analysis’. In: Briefings in Bioinformatics 22.3 (2021), bbaa148. ISSN: 1477-4054. DOI: 10.1093/bib/bbaa148.

[20] Ryan Poplin et al. ‘A universal SNP and small-indel variant caller using deep neural networks’. In: Nature Biotechnology 36.10 (2018), pp. 983–987. ISSN: 1087-0156. DOI: 10.1038/nbt.4235.

[21] Alan Tourancheau et al. ‘Discovering multiple types of DNA methylation from bacteria and microbiome using nanopore sequencing’. In: Nature methods 18.5 (2021), pp. 491–498. ISSN: 1548-7091. DOI: 10.1038/s41592-021-01109-3.

[22] Stephen J Bush. ‘Generalizable characteristics of false-positive bacterial variant calls’. In: Microbial Genomics 7.8 (2021). DOI: 10.1099/mgen.0.000615.

[23] Stephen J Bush et al. ‘Genomic diversity affects the accuracy of bacterial single-nucleotide polymorphism–calling pipelines’. In: GigaScience 9.2 (2020), giaa007.#x2013;. ISSN: 2047-217X. DOI: 10.1093/gigascience/giaa007.

[24] Sina Majidian et al. ‘Genomic variant benchmark: if you cannot measure it, you cannot improve it’. In: Genome Biology 24.1 (2023), p. 221. ISSN: 1474-760X. DOI: 10.1186/s13059-023-03061-1.

[25] Heng Li. ‘Toward better understanding of artifacts in variant calling from high-coverage samples’. In: Bioinformatics 30.20 (2014), pp. 2843–2851. ISSN: 1367-4803. DOI: 10.1093/bioinformatics/btu356.

[26] Heng Li et al. ‘A synthetic-diploid benchmark for accurate variant-calling evaluation’. In: Nature Methods 15.8 (2018), pp. 595–597. ISSN: 1548-7105. DOI: 10.1038/s41592-018-0054-7.

[27] Heng Li. ‘Minimap2: pairwise alignment for nucleotide sequences’. In: Bioinformatics 34.18 (2018), pp. 3094–3100. ISSN: 1367-4803. DOI: 10.1093/bioinformatics/bty191.

[28] Guillaume Marçais et al. ‘MUMmer4: A fast and versatile genome alignment system’. In: PLOS Computational Biology 14.1 (2018), e1005944. ISSN: 1553-734X. DOI: 10.1371/journal.pcbi.1005944.

[29] Petr Danecek et al. ‘Twelve years of SAMtools and BCFtools’. In: GigaScience 10.2 (2021), giab008. ISSN: 2047-217X. DOI: 10.1093/gigascience/giab008.

[30] Erik Garrison and Gabor Marth. Haplotype-based variant detection from short-read sequencing. 2012. DOI: 10.48550/arXiv.1207.3907.

[31] Oxford Nanopore Technologies. Medaka: Sequence correction provided by ONT Research. 2023. URL: https://github.com/nanoporetech/medaka.

[32] Torsten Seemann. snippy: fast bacterial variant calling from NGS reads. Version 4.6.0. 2015. URL: https://github.com/tseemann/snippy.

[33] Tim Dunn and Satish Narayanasamy. ‘vcfdist: accurately benchmarking phased small variant calls in human genomes’. In: Nature Communications 14.1 (2023), p. 8149. ISSN: 2041-1723. DOI: 10.1038/s41467-023-43876-x.

[34] Todd J. Treangen and Steven L. Salzberg. ‘Repetitive DNA and next-generation sequencing: computational challenges and solutions’. In: Nature Reviews Genetics 13.1 (2012), pp. 36–46. ISSN: 1471-0064. DOI: 10.1038/nrg3117.

[35] Michael B. Hall. ‘Rasusa: Randomly subsample sequencing reads to a specified coverage’. In: Journal of Open Source Software 7.69 (2022), p. 3941. ISSN: 2475-9066. DOI: 10.21105/joss.03941.

[36] Ryan R. Wick, Louise M. Judd, and Kathryn E. Holt. ‘Assembling the perfect bacterial genome using Oxford Nanopore and Illumina sequencing’. In: PLOS Computational Biology 19.3 (2023), e1010905. ISSN: 1553-7358. DOI: 10.1371/journal.pcbi.1010905.

[37] Teresa L. Street et al. ‘Optimizing DNA Extraction Methods for Nanopore Sequencing of Neisseria gonorrhoeae Directly from Urine Samples’. In: Journal of Clinical Microbiology 58.3 (2020), 10.1128/jcm.01822–19. DOI: 10.1128/jcm.01822-19.

[38] Charles Y. Chiu and Steven A. Miller. ‘Clinical metagenomics’. In: Nature Reviews Genetics 20.6 (2019), pp. 341–355. ISSN: 1471-0064. DOI: 10.1038/s41576-019-0113-7.

[39] Kayzad Nilgiriwala et al. ‘Genomic Sequencing from Sputum for Tuberculosis Disease Diagnosis, Lineage Determination, and Drug Susceptibility Prediction’. In: Journal of Clinical Microbiology 61.3 (2023), e01578–22. DOI: 10.1128/jcm.01578-22.

[40] Lillian Musila. ‘Genomic outbreak surveillance in resource-poor settings’. In: Nature Reviews Genetics (2022), pp. 1–1. ISSN: 1471-0056. DOI: 10.1038/s41576-022-00500-w.

[41] Anbo Zhou, Timothy Lin, and Jinchuan Xing. ‘Evaluating nanopore sequencing data processing pipelines for structural variation identification’. In: Genome Biology 20.1 (2019), p. 237. ISSN: 1474-760X. DOI: 10.1186/s13059-019-1858-1.

[42] Nicholas Stoler and Anton Nekrutenko. ‘Sequencing error profiles of Illumina sequencing instruments’. In: NAR Genomics and Bioinformatics 3.1 (2021), lqab019. ISSN: 2631-9268. DOI: 10.1093/nargab/lqab019.

[43] Wei Shen et al. ‘SeqKit: A Cross-Platform and Ultrafast Toolkit for FASTA/Q File Manipulation’. In: PLOS ONE 11.10 (2016), e0163962. ISSN: 1932-6203. DOI: 10.1371/journal.pone.0163962.

[44] Shifu Chen et al. ‘fastp: an ultra-fast all-in-one FASTQ preprocessor’. In: Bioinformatics 34.17 (2018), pp. i884–i890. ISSN: 1367-4803. DOI: 10.1093/bioinformatics/bty560.

[45] Ryan R. Wick. Filtlong: quality filtering tool for long reads. Version 0.2.1. 2021. URL: https://github.com/rrwick/Filtlong.

[46] Ryan R. Wick et al. ‘Trycycler: consensus long-read assemblies for bacterial genomes’. In: Genome Biology 22.1 (2021), p. 266. ISSN: 1474-760X. DOI: 10.1186/s13059-021-02483-z.

[47] Ryan R. Wick and Kathryn E. Holt. ‘Polypolish: Short-read polishing of long-read bacterial genome assemblies’. In: PLOS Computational Biology 18.1 (2022), e1009802. ISSN: 1553-7358. DOI: 10.1371/journal.pcbi.1009802.

[48] George Bouras et al. ‘How low can you go? Short-read polishing of Oxford Nanopore bacterial genome assemblies’. In: bioRxiv (2024). DOI: 10.1101/2024.03.07.584013.

[49] Aleksey V. Zimin and Steven L. Salzberg. ‘The genome polishing tool POLCA makes fast and accurate corrections in genome assemblies’. In: PLOS Computational Biology 16.6 (2020), e1007981. ISSN: 1553-7358. DOI: 10.1371/journal.pcbi.1007981.

[50] Vitor C. Piro. genome_updater. Version 0.6.3. 2023. DOI: 10.5281/zenodo.8108640.

[51] Jim Shaw and Yun William Yu. ‘Fast and robust metagenomic sequence comparison through sparse chaining with skani’. In: Nature Methods 20.11 (2023), pp. 1661–1665. ISSN: 1548-7105. DOI: 10.1038/s41592-023-02018-3.

[52] Donovan H. Parks et al. ‘CheckM: assessing the quality of microbial genomes recovered from isolates, single cells, and metagenomes’. In: Genome Research 25.7 (2015), pp. 1043–1055. ISSN: 1088-9051, 1549-5469. DOI: 10.1101/gr.186072.114.

[53] Luis M. Rodriguez-R et al. ‘An ANI gap within bacterial species that advances the definitions of intra-species units’. In: mBio 15.1 (2023), e02696–23. DOI: 10.1128/mbio.02696-23.

[54] Tomeu Viver et al. ‘Towards estimating the number of strains that make up a natural bacterial population’. In: Nature Communications 15.1 (2024), p. 544. ISSN: 2041-1723. DOI: 10.1038/s41467-023-44622-z.

[55] Michael B. Hall. mbhall88/NanoVarBench. 2024. DOI: 10.5281/zenodo.10820970.

[56] Michael B. Hall. NanoVarBench: Evaluating Nanopore-based bacterial variant calling. 2023. URL: https://github.com/mbhall88/NanoVarBench.

[57] Oxford Nanopore Technologies. Rerio: Research release basecalling models and configurations. 2023. URL: https://github.com/nanoporetech/rerio.

[58] Heng Li. ‘Aligning sequence reads, clone sequences and assembly contigs with BWA-MEM’. In: arXiv (2013). DOI: 10.48550/arXiv.1303.3997.

[59] Aaron R. Quinlan and Ira M. Hall. ‘BEDTools: a flexible suite of utilities for comparing genomic features’. In: Bioinformatics 26.6 (2010), pp. 841–842. ISSN: 1367-4803. DOI: 10.1093/bioinformatics/btq033.

[60] Michael B. Hall. NanoVarBench variant truthset files. Zenodo, 2024. DOI: 10.5281/zenodo.10867171.

