## Supplementary Material for "Benchmarking reveals superiority of deep learning variant callers on bacterial nanopore sequence data"

#### Contents

**S1 Sample selection** **6**

**S2 Simplex F1 outlier** **6**

**References** **28**

#### List of Figures

S1 Pileup surrounding a Clair3 false negative variant. The top track (four blue rectangles) shows true variants - i.e., variants we expect to find. The second track shows a single false negative variant. The third track shows the alignment of ONT reads to this region, with coloured letters indicating where the aligned sequence disagrees with the reference. The fourth track shows the alignment of Illumina reads. The fifth track is the reference sequence. The bottom track shows repetitive regions (the whole region is repetitive as the blue rectangle spans the whole region in view). 7

S2 Textual example of two small repetitive regions in sample ATCC_25922 that lead to a false negative at position 3 in sequence S1. Positions 1 and 2 are two other variant positions where a true positive was obtained for S1 (the expected variants at positions 1 and 2 match the sequence in S2). . . . . . . . . . . . . . . . . . . . . . . . 7

S3 Precision at the highest F1 score for each sample (points), stratified by basecalling model (colours), variant type (rows), and read type (columns). Illumina results (green) are included as a reference and do no have different basecalling models or read types. Note, longshot does not provide indel calls. . . . . . . . . . . . . 8

S4 Recall at the highest F1 score for each sample (points), stratified by basecalling model (colours), variant type (rows), and read type (columns). Illumina results (green) are included as a reference and do no have different basecalling models or read types. Note, longshot does not provide indel calls. . . . . . . . . . . . . . . . . 9
S5 Clair3 sup model F1 score (y-axis) at the highest F1 score for each sample (x-axis), stratified by variant type (rows), and read type (shapes). . . . . . . . . . . . . . . . . . . . . . . . . . . . . . 10
S6 Clair3 sup model precision (y-axis) at the highest F1 score for each sample (x-axis), stratified by variant type (rows), and read type (shapes). . . . . . . . . . . . . . . . . . . . . . . . . . . . . . 11
S7 Clair3 sup model recall (y-axis) at the highest F1 score for each sample (x-axis), stratified by variant type (rows), and read type (shapes). . . . . . . . . . . . . . . . . . . . . . . . . . . . . . . 12
S8 Precision and recall curves for each variant caller (colours and line styles) on sequencing data basecalled with the hac model, stratified by variant type (rows) and ready type (columns). The curves are generated by using increasing variant quality score thresholds to filter variants and calculating precision and recall at each threshold. The lowest threshold is the lower right part of the curve, moving to the highest at the top left. Note, Longshot does not provide indel calls. . . . . . . . . . . . . . . . . . . . . 13
S9 Precision and recall curves for each variant caller (colours and line styles) on sequencing data basecalled with the fast model, stratified by variant type (rows). The curves are generated by using increasing variant quality score thresholds to filter variants and calculating precision and recall at each threshold. The lowest threshold is the lower right part of the curve, moving to the highest at the top left. Note, longshot does not provide indel calls. 14
S10 Read pileup in a variant-dense region in the genome of sample ATCC_17802__202309. The top track shows the alignment of the sample's Illumina reads, while the lower track is the ONT reads. Missed variant calls (false negatives) are shown by small blue notches at the bottom of the figure, but are also identifiable by vertical coloured lines in the ONT reads. Pileup is visualised in IGV [1]. . . . . . . . . . . . . . . . . . . . . . . . . . . . . . . 15
S11 Read pileup around two repetitive regions (horizontal blue bars) in the genome of sample AMtb_1__202401. The top track shows the alignment of the sample's Illumina reads, while the lower track is the ONT reads. Missed variant calls (false negatives) are shown by small red notches in a purple box with a red border at the bottom of the figure, but are also identifiable by vertical coloured lines in the ONT reads. Pileup is visualised in IGV [1]. 15

S12 Relationship between indel length (x-axis) and homopolymer length (y-axis) for false positive (FP) indel calls for BCFtools 100x simplex fast (top left), hac (top right), and sup (lower left) calls. Illumina is shown in the lower right for reference. The vertical red line indicates the threshold above which we deem a run of the same nucleotide to be a "true" homopolymer. Indel length is the number of bases inserted/deleted for an indel, whereas the homopolymer length indicates how long the tract of the same nucleotide is after the indel. The colour of a cell indicates how many FP indels of that indel-homopolymer length combination. . 16
S13 Relationship between indel length (x-axis) and homopolymer length (y-axis) for false positive (FP) indel calls for DeepVariant 100x simplex fast (top left), hac (top right), and sup (lower left) calls. Illumina is shown in the lower right for reference. The vertical red line indicates the threshold above which we deem a run of the same nucleotide to be a "true" homopolymer. Indel length is the number of bases inserted/deleted for an indel, whereas the homopolymer length indicates how long the tract of the same nucleotide is after the indel. The colour of a cell indicates how many FP indels of that indel-homopolymer length combination. . 17
S14 Relationship between indel length (x-axis) and homopolymer length (y-axis) for false positive (FP) indel calls for FreeBayes 100x simplex fast (top left), hac (top right), and sup (lower left) calls. Illumina is shown in the lower right for reference. The vertical red line indicates the threshold above which we deem a run of the same nucleotide to be a "true" homopolymer. Indel length is the number of bases inserted/deleted for an indel, whereas the homopolymer length indicates how long the tract of the same nucleotide is after the indel. The colour of a cell indicates how many FP indels of that indel-homopolymer length combination. . 18
S15 Relationship between indel length (x-axis) and homopolymer length (y-axis) for false positive (FP) indel calls for Medaka 100x simplex fast (top left), hac (top right), and sup (lower left) calls. Illumina is shown in the lower right for reference. The vertical red line indicates the threshold above which we deem a run of the same nucleotide to be a "true" homopolymer. Indel length is the number of bases inserted/deleted for an indel, whereas the homopolymer length indicates how long the tract of the same nucleotide is after the indel. The colour of a cell indicates how many FP indels of that indel-homopolymer length combination. . 19

S16 Relationship between indel length (x-axis) and homopolymer length (y-axis) for false positive (FP) indel calls for NanoCaller 100x simplex fast (top left), hac (top right), and sup (lower left) calls. Illumina is shown in the lower right for reference. The vertical red line indicates the threshold above which we deem a run of the same nucleotide to be a “true” homopolymer. Indel length is the number of bases inserted/deleted for an indel, whereas the homopolymer length indicates how long the tract of the same nucleotide is after the indel. The colour of a cell indicates how many FP indels of that indel-homopolymer length combination. . 20
S17 Relationship between indel length (x-axis) and homopolymer length (y-axis) for false negative (FN) indel calls for Clair3 100x simplex fast (top left), hac (top right), and sup (lower left) calls. Illumina is shown in the lower right for reference. The vertical red line indicates the threshold above which we deem a run of the same nucleotide to be a “true” homopolymer. Indel length is the number of bases inserted/deleted for an indel, whereas the homopolymer length indicates how long the tract of the same nucleotide is after the indel. The colour of a cell indicates how many FN indels of that indel-homopolymer length combination. . 21
S18 Relationship between indel length (x-axis) and homopolymer length (y-axis) for false negative (FN) indel calls for BCFtools 100x simplex fast (top left), hac (top right), and sup (lower left) calls. Illumina is shown in the lower right for reference. The vertical red line indicates the threshold above which we deem a run of the same nucleotide to be a “true” homopolymer. Indel length is the number of bases inserted/deleted for an indel, whereas the homopolymer length indicates how long the tract of the same nucleotide is after the indel. The colour of a cell indicates how many FN indels of that indel-homopolymer length combination. . 22
S19 Relationship between indel length (x-axis) and homopolymer length (y-axis) for false negative (FN) indel calls for DeepVariant 100x simplex fast (top left), hac (top right), and sup (lower left) calls. Illumina is shown in the lower right for reference. The vertical red line indicates the threshold above which we deem a run of the same nucleotide to be a “true” homopolymer. Indel length is the number of bases inserted/deleted for an indel, whereas the homopolymer length indicates how long the tract of the same nucleotide is after the indel. The colour of a cell indicates how many FN indels of that indel-homopolymer length combination. . 23

S20 Relationship between indel length (x-axis) and homopolymer length (y-axis) for false negative (FN) indel calls for FreeBayes 100x simplex fast (top left), hac (top right), and sup (lower left) calls. Illumina is shown in the lower right for reference. The vertical red line indicates the threshold above which we deem a run of the same nucleotide to be a “true” homopolymer. Indel length is the number of bases inserted/deleted for an indel, whereas the homopolymer length indicates how long the tract of the same nucleotide is after the indel. The colour of a cell indicates how many FN indels of that indel-homopolymer length combination. . 24
S21 Relationship between indel length (x-axis) and homopolymer length (y-axis) for false negative (FN) indel calls for Medaka 100x simplex fast (top left), hac (top right), and sup (lower left) calls. Illumina is shown in the lower right for reference. The vertical red line indicates the threshold above which we deem a run of the same nucleotide to be a “true” homopolymer. Indel length is the number of bases inserted/deleted for an indel, whereas the homopolymer length indicates how long the tract of the same nucleotide is after the indel. The colour of a cell indicates how many FN indels of that indel-homopolymer length combination. . 25
S22 Relationship between indel length (x-axis) and homopolymer length (y-axis) for false negative (FN) indel calls for NanoCaller 100x simplex fast (top left), hac (top right), and sup (lower left) calls. Illumina is shown in the lower right for reference. The vertical red line indicates the threshold above which we deem a run of the same nucleotide to be a “true” homopolymer. Indel length is the number of bases inserted/deleted for an indel, whereas the homopolymer length indicates how long the tract of the same nucleotide is after the indel. The colour of a cell indicates how many FN indels of that indel-homopolymer length combination. . 26
S23 Runtime of basecalling ONT data with different models on GPUs. Runtime is presented as seconds per megabasepairs, keeping consistent with Figure 8 from the main text. The y-axis represents basecalling with either 8 Nvidia A100 GPUs (cuda:all) or 1 A100 (cuda:single), with each stratified by basecalling model (colours). Points represent running a single sample-model combination, with each combination being run three times. . . . . . . 27

#### Abbreviations

- FN: false negative
- ONT: Oxford Nanopore Technologies

### S1 Sample selection

MMC234__202311 and RDH275__202311 were collected as part of an invasive streptococcal surveillance study in Australia (approved by the Royal Melbourne Hospital Human Research Ethics Committee [HREC/80105/MH-2021] and the Human Research Ethics Committee of the Northern Territory Depart of Health and Menzies School of Health Research [2021-4181]).

### S2 Simplex F1 outlier

From Figure 2 in the main text it is clear that in the simplex SNP panel (top left) that there is a single outlier for the sup model for most variant callers. This sample is the same across the variant callers - ATCC_25922 (*E. coli*). We chose to investigate the reason for this outlier using the Clair3 (simplex sup) variant calls. The reason for the lower F1 score was reduced recall (ass seen in Figure S4). ATCC_25922 had 47 false negative (FN) variants, with 45 of those coming from two repetitive regions (see *Identifying repetitive regions* in the main text). In addition, these two regions have a higher variant density compared to the rest of the genome and are the repeats of each other. The two regions are 7.5Kbp long and have an identity of 96.5%. As can be seen from Figure S1, there is a lot of heterogeneity at the FN site. This is caused by reads from the reciprocal region multi-mapping to this region. Figure S2 shows a textual representation of the of the two sequences. Position 3 is the FN, where we expect a C¿T variant call. While 50% of the bases at position 3 are indeed T, and 34% are C, Clair3 and most other callers fail to call this site a variant site. This pattern of heterogeneity leading to missed calls was repeated for the other FNs within these repeats. Highlighting that even though deep learning callers using R10 ONT data deal much better with repetitive and variant dense regions, they can still struggle when you have both of these difficulties combined to a high degree. However, it is interesting to note that Medaka did not have the same issues with missed variants for these sites. Assumably the model training for Medaka is sufficiently different to DeepVariant and Clair3 to allow it to deal with these difficult sites without issue.

The other notable outlier from Figure 2 in the main text is in the duplex SNP panel (top right). This outlier is sample KPC2_202310 (*K. pneumoniae*). The reason for its reduced F1 score is again to do with recall. However, this reduced recall is simply due to the fact that the duplex depth for this sample is only 3x.

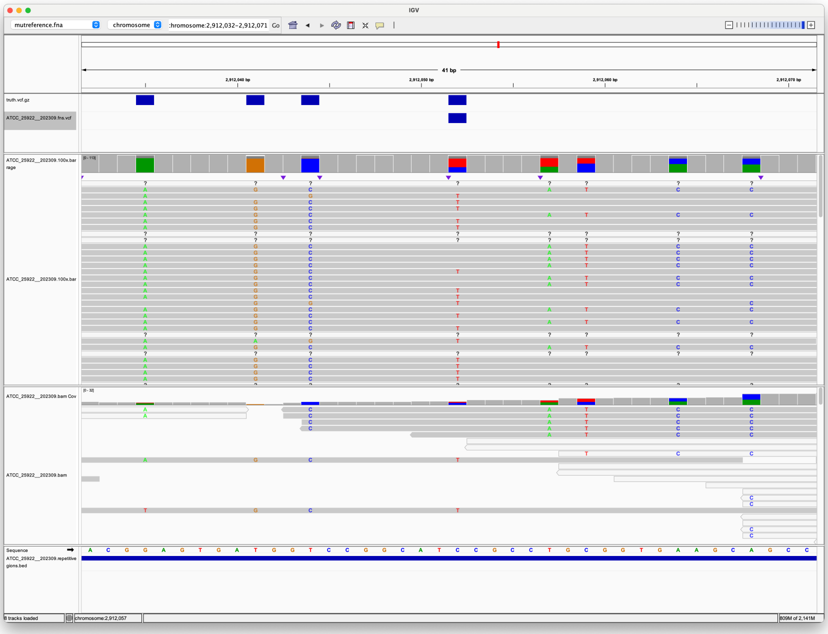

**Figure S1:** Pileup surrounding a Clair3 false negative variant. The top track (four blue rectangles) shows true variants - i.e., variants we expect to find. The second track shows a single false negative variant. The third track shows the alignment of ONT reads to this region, with coloured letters indicating where the aligned sequence disagrees with the reference. The fourth track shows the alignment of Illumina reads. The fifth track is the reference sequence. The bottom track shows repetitive regions (the whole region is repetitive as the blue rectangle spans the whole region in view).

```
        1     2          3
S1 - {T}GG{T}CCGGCAT{C}CGCC
S2 - {G}GG{C}CCGGCAT{T}CGCC
        1     2          3
```

**Figure S2:** Textual example of two small repetitive regions in sample ATCC_25922 that lead to a false negative at position 3 in sequence S1. Positions 1 and 2 are two other variant positions where a true positive was obtained for S1 (the expected variants at positions 1 and 2 match the sequence in S2).

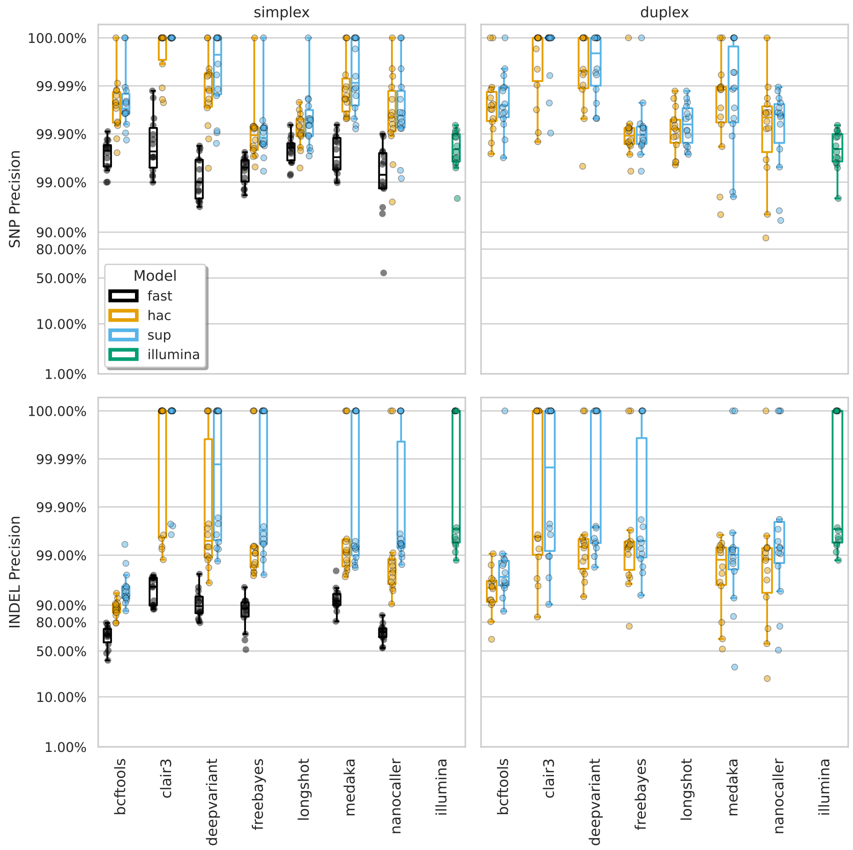


**Figure S3:** Precision at the highest F1 score for each sample (points), stratified by basecalling model (colours), variant type (rows), and read type (columns). Illumina results (green) are included as a reference and do no have different basecalling models or read types. Note, longshot does not provide indel calls.

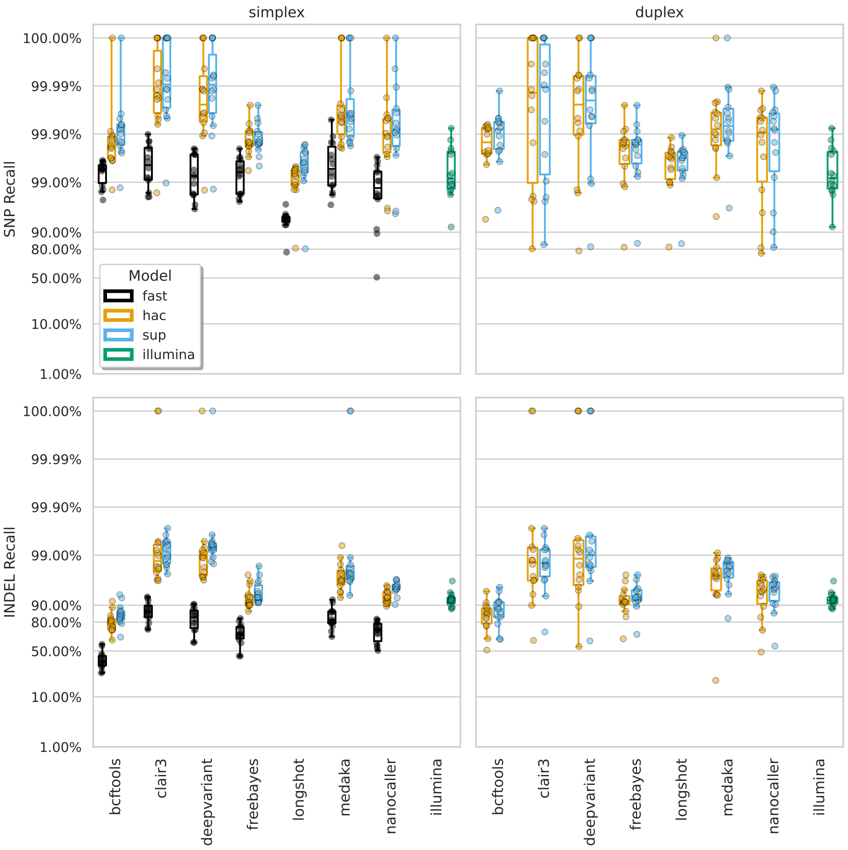


**Figure S4:** Recall at the highest F1 score for each sample (points), stratified by basecalling model (colours), variant type (rows), and read type (columns). Illumina results (green) are included as a reference and do no have different basecalling models or read types. Note, longshot does not provide indel calls.

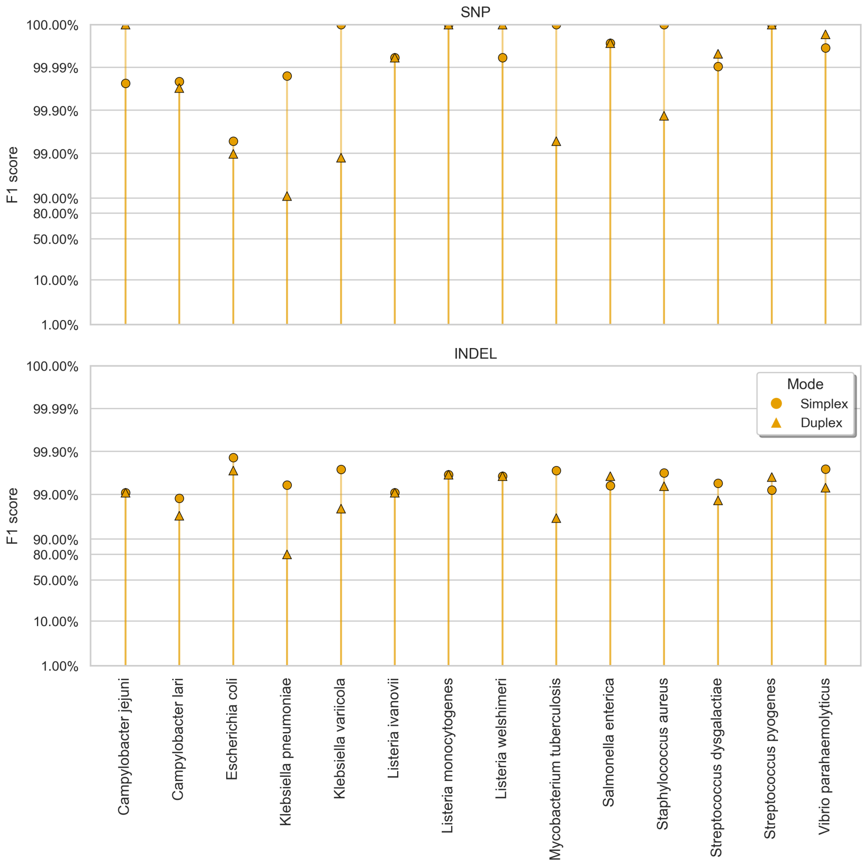


**Figure S5:** Clair3 sup model F1 score (y-axis) at the highest F1 score for each sample (x-axis), stratified by variant type (rows), and read type (shapes).

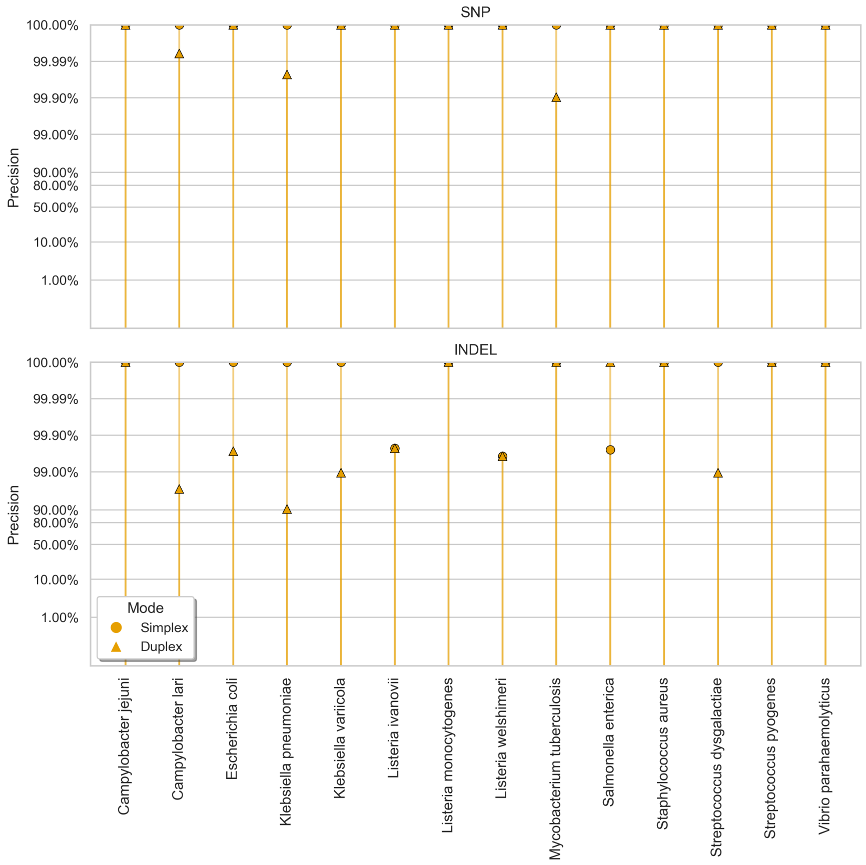


**Figure S6:** Clair3 sup model precision (y-axis) at the highest F1 score for each sample (x-axis), stratified by variant type (rows), and read type (shapes).

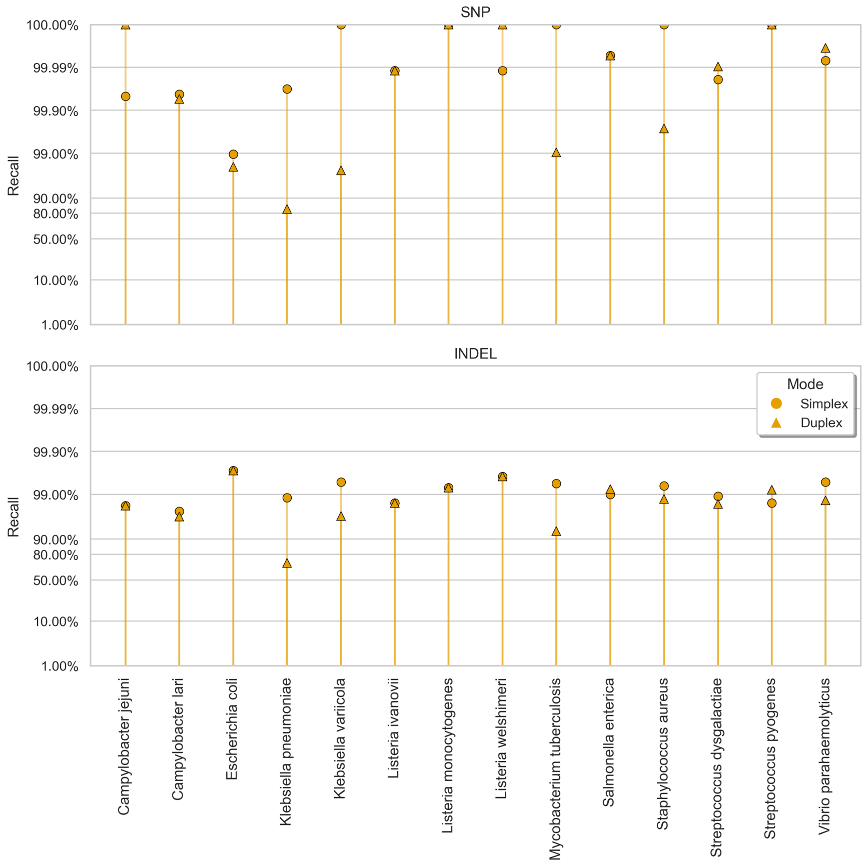


**Figure S7:** Clair3 sup model recall (y-axis) at the highest F1 score for each sample (x-axis), stratified by variant type (rows), and read type (shapes).

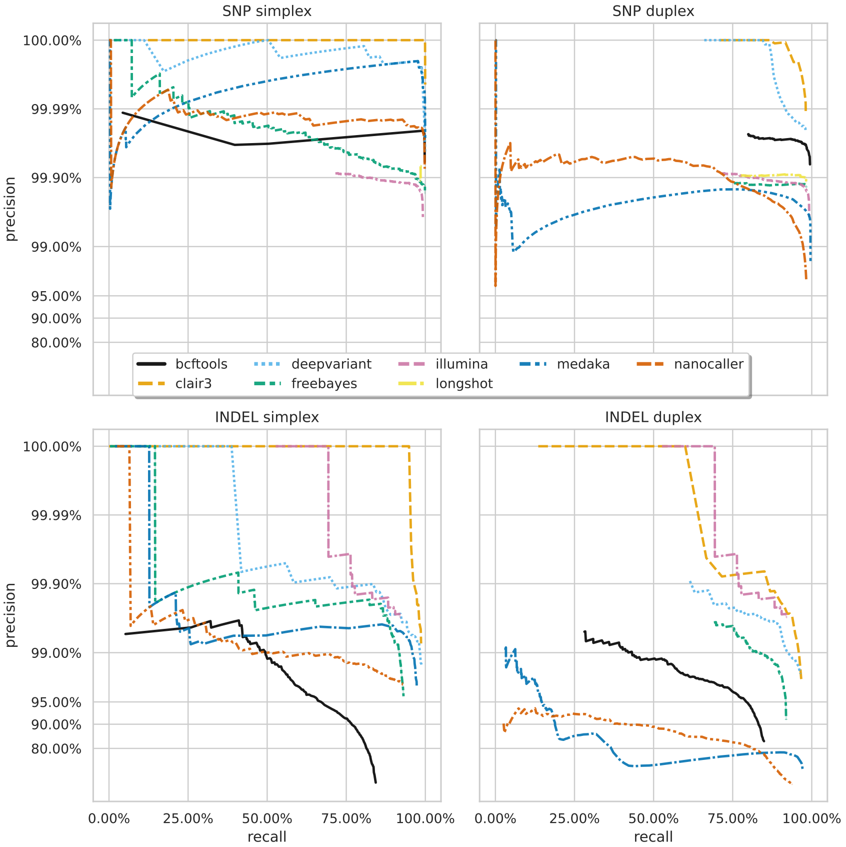


**Figure S8:** Precision and recall curves for each variant caller (colours and line styles) on sequencing data basecalled with the hac model, stratified by variant type (rows) and ready type (columns). The curves are generated by using increasing variant quality score thresholds to filter variants and calculating precision and recall at each threshold. The lowest threshold is the lower right part of the curve, moving to the highest at the top left. Note, Longshot does not provide indel calls.

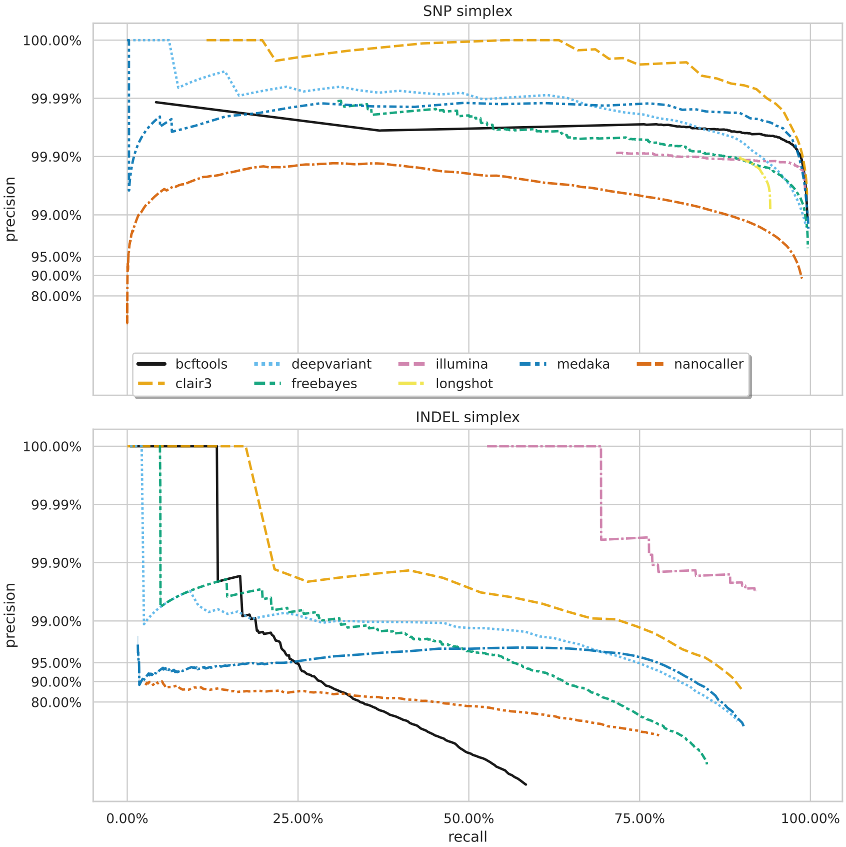


**Figure S9:** Precision and recall curves for each variant caller (colours and line styles) on sequencing data basecalled with the fast model, stratified by variant type (rows). The curves are generated by using increasing variant quality score thresholds to filter variants and calculating precision and recall at each threshold. The lowest threshold is the lower right part of the curve, moving to the highest at the top left. Note, longshot does not provide indel calls.

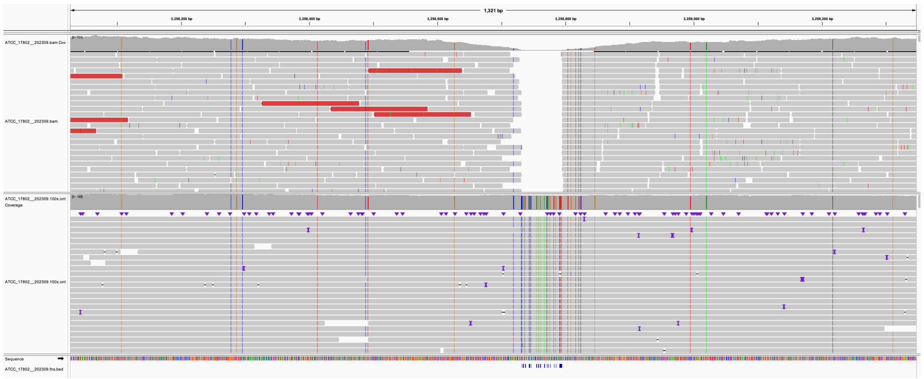

**Figure S10:** Read pileup in a variant-dense region in the genome of sample ATCC_17802__202309. The top track shows the alignment of the sample's Illumina reads, while the lower track is the ONT reads. Missed variant calls (false negatives) are shown by small blue notches at the bottom of the figure, but are also identifiable by vertical coloured lines in the ONT reads. Pileup is visualised in IGV [1].

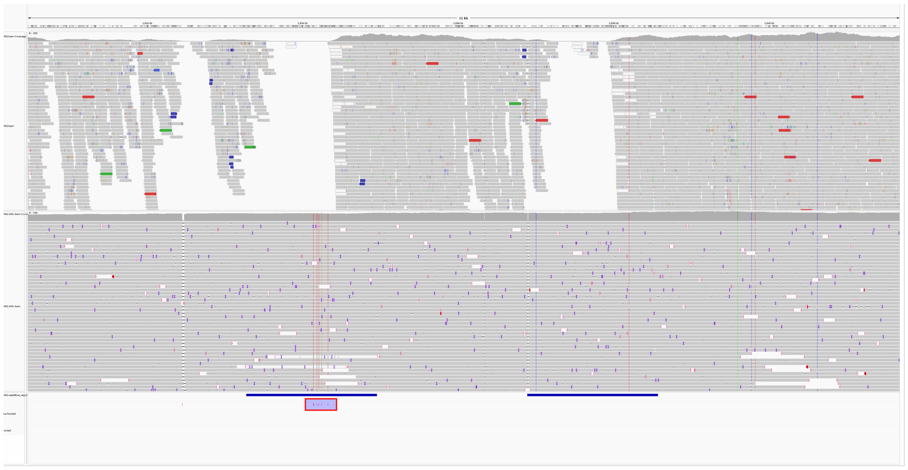

**Figure S11:** Read pileup around two repetitive regions (horizontal blue bars) in the genome of sample AMtb_1__202401. The top track shows the alignment of the sample's Illumina reads, while the lower track is the ONT reads. Missed variant calls (false negatives) are shown by small red notches in a purple box with a red border at the bottom of the figure, but are also identifiable by vertical coloured lines in the ONT reads. Pileup is visualised in IGV [1].

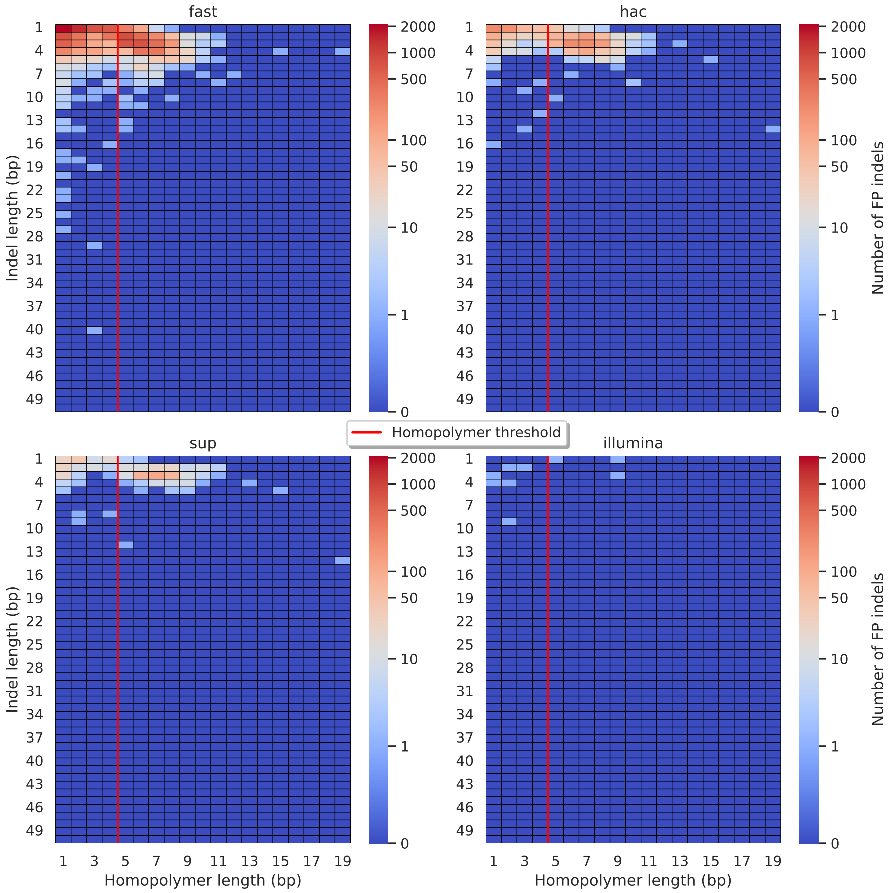


**Figure S12:** Relationship between indel length (x-axis) and homopolymer length (y-axis) for false positive (FP) indel calls for BCFtools 100x simplex fast (top left), hac (top right), and sup (lower left) calls. Illumina is shown in the lower right for reference. The vertical red line indicates the threshold above which we deem a run of the same nucleotide to be a “true” homopolymer. Indel length is the number of bases inserted/deleted for an indel, whereas the homopolymer length indicates how long the tract of the same nucleotide is after the indel. The colour of a cell indicates how many FP indels of that indel-homopolymer length combination.

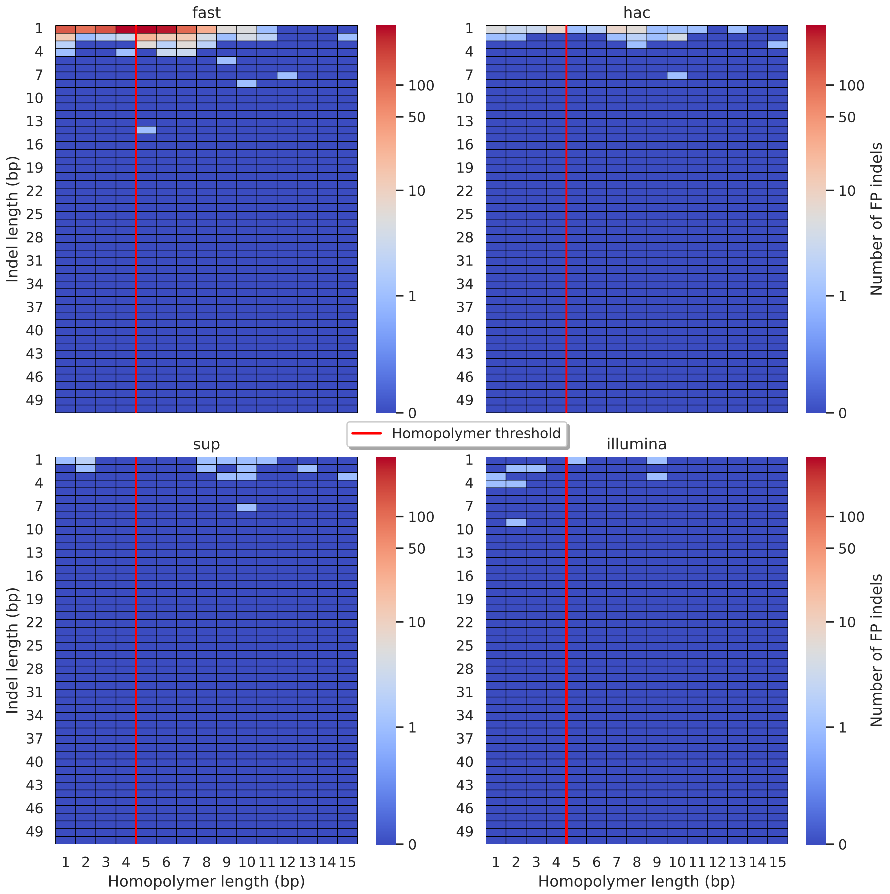


**Figure S13:** Relationship between indel length (x-axis) and homopolymer length (y-axis) for false positive (FP) indel calls for DeepVariant 100x simplex fast (top left), hac (top right), and sup (lower left) calls. Illumina is shown in the lower right for reference. The vertical red line indicates the threshold above which we deem a run of the same nucleotide to be a "true" homopolymer. Indel length is the number of bases inserted/deleted for an indel, whereas the homopolymer length indicates how long the tract of the same nucleotide is after the indel. The colour of a cell indicates how many FP indels of that indel-homopolymer length combination.

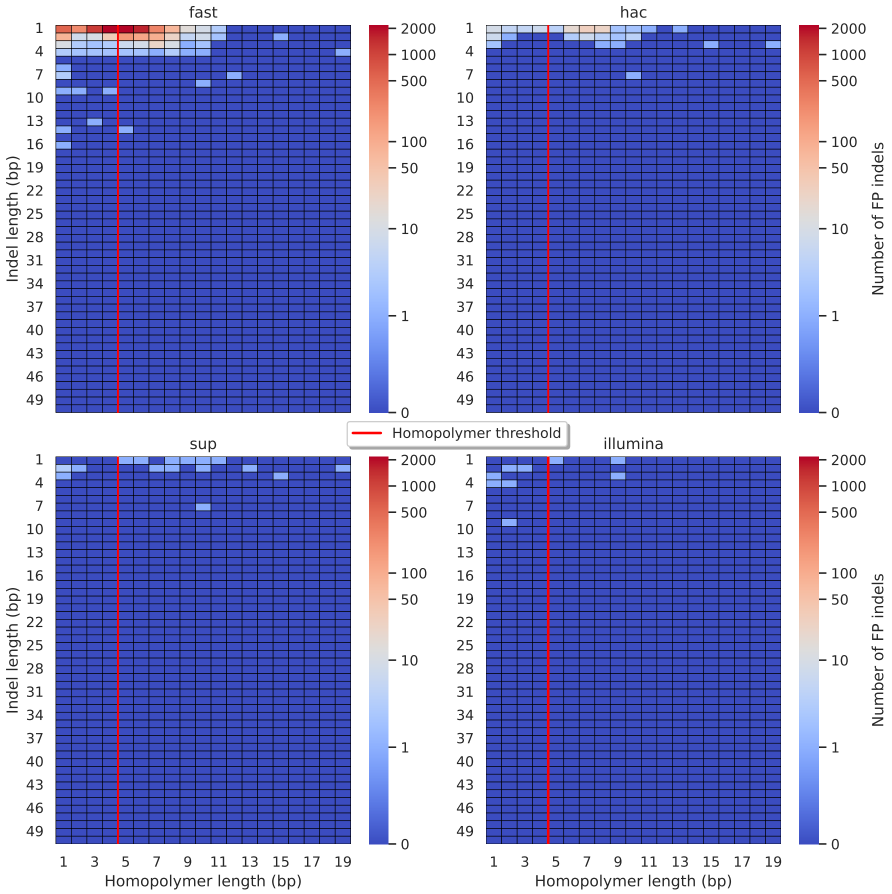


**Figure S14:** Relationship between indel length (x-axis) and homopolymer length (y-axis) for false positive (FP) indel calls for FreeBayes 100x simplex fast (top left), hac (top right), and sup (lower left) calls. Illumina is shown in the lower right for reference. The vertical red line indicates the threshold above which we deem a run of the same nucleotide to be a "true" homopolymer. Indel length is the number of bases inserted/deleted for an indel, whereas the homopolymer length indicates how long the tract of the same nucleotide is after the indel. The colour of a cell indicates how many FP indels of that indel-homopolymer length combination.

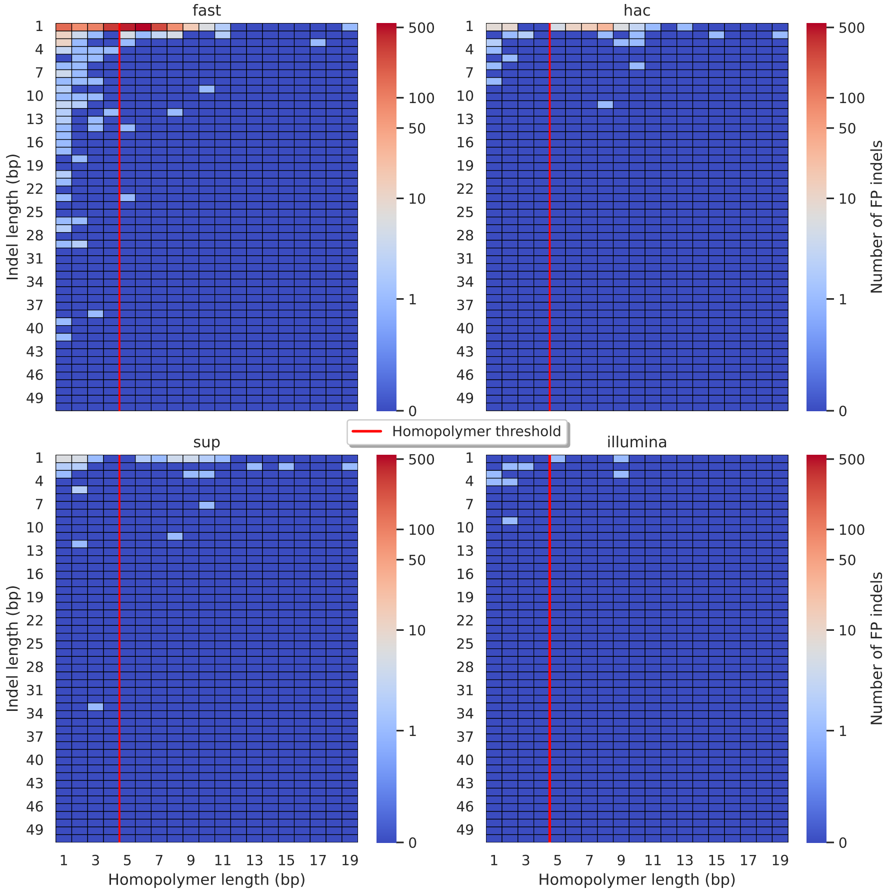


**Figure S15:** Relationship between indel length (x-axis) and homopolymer length (y-axis) for false positive (FP) indel calls for Medaka 100x simplex fast (top left), hac (top right), and sup (lower left) calls. Illumina is shown in the lower right for reference. The vertical red line indicates the threshold above which we deem a run of the same nucleotide to be a "true" homopolymer. Indel length is the number of bases inserted/deleted for an indel, whereas the homopolymer length indicates how long the tract of the same nucleotide is after the indel. The colour of a cell indicates how many FP indels of that indel-homopolymer length combination.

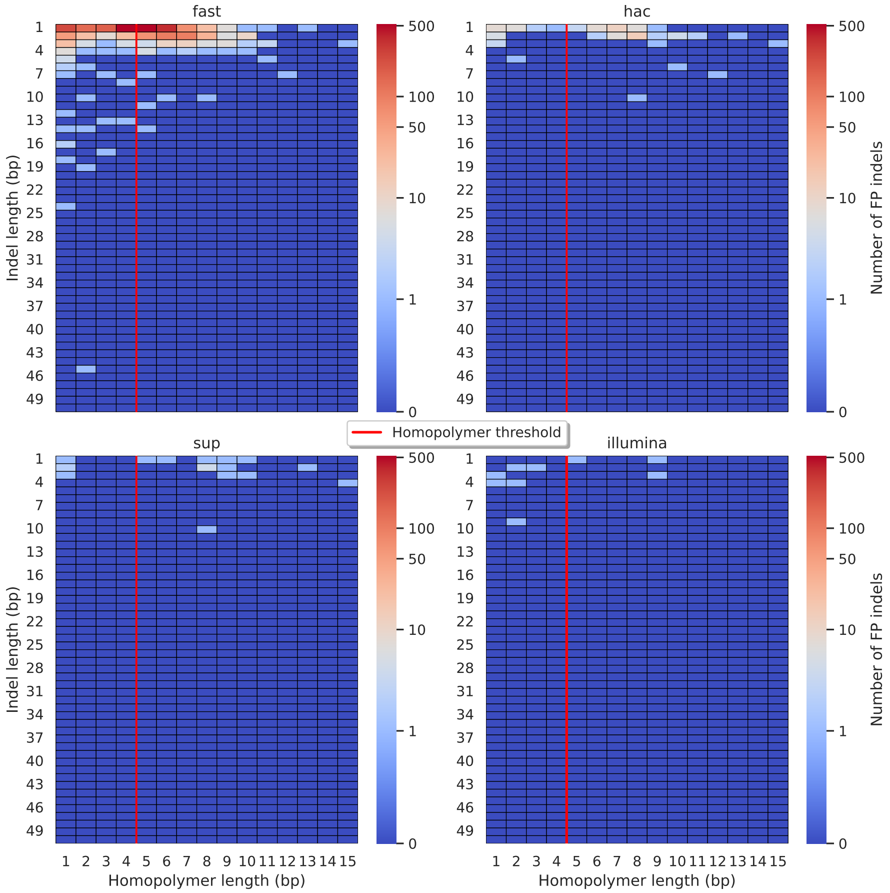


**Figure S16:** Relationship between indel length (x-axis) and homopolymer length (y-axis) for false positive (FP) indel calls for NanoCaller 100x simplex fast (top left), hac (top right), and sup (lower left) calls. Illumina is shown in the lower right for reference. The vertical red line indicates the threshold above which we deem a run of the same nucleotide to be a “true” homopolymer. Indel length is the number of bases inserted/deleted for an indel, whereas the homopolymer length indicates how long the tract of the same nucleotide is after the indel. The colour of a cell indicates how many FP indels of that indel-homopolymer length combination.

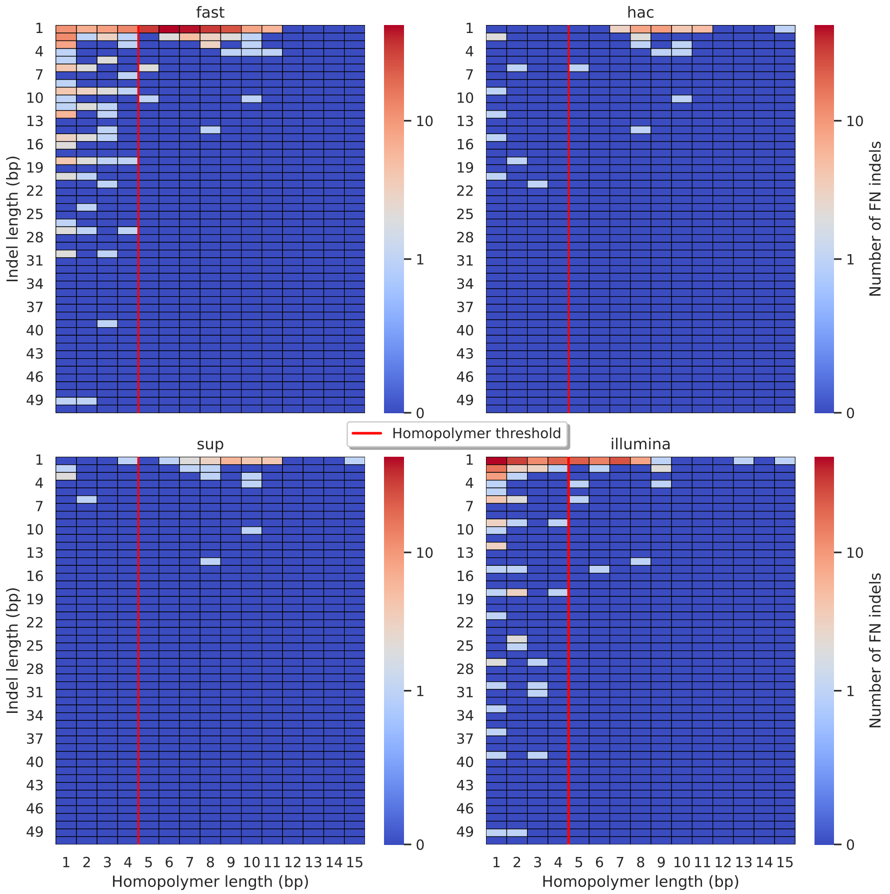


**Figure S17:** Relationship between indel length (x-axis) and homopolymer length (y-axis) for false negative (FN) indel calls for Clair3 100x simplex fast (top left), hac (top right), and sup (lower left) calls. Illumina is shown in the lower right for reference. The vertical red line indicates the threshold above which we deem a run of the same nucleotide to be a "true" homopolymer. Indel length is the number of bases inserted/deleted for an indel, whereas the homopolymer length indicates how long the tract of the same nucleotide is after the indel. The colour of a cell indicates how many FN indels of that indel-homopolymer length combination.

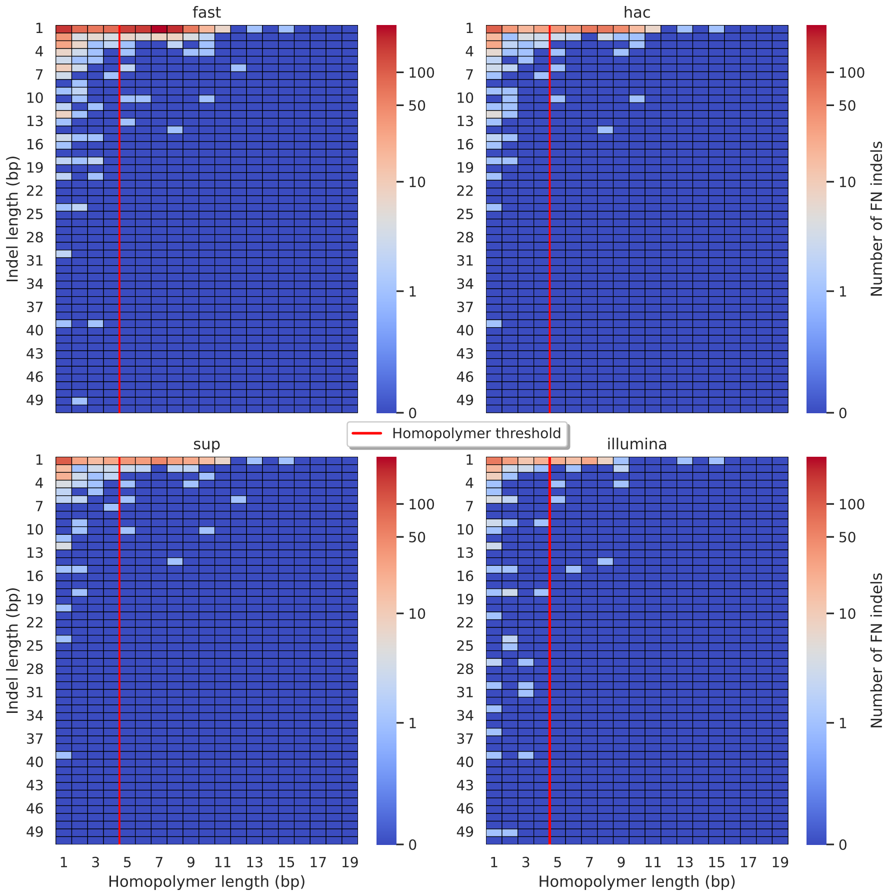


**Figure S18:** Relationship between indel length (x-axis) and homopolymer length (y-axis) for false negative (FN) indel calls for BCFtools 100x simplex fast (top left), hac (top right), and sup (lower left) calls. Illumina is shown in the lower right for reference. The vertical red line indicates the threshold above which we deem a run of the same nucleotide to be a "true" homopolymer. Indel length is the number of bases inserted/deleted for an indel, whereas the homopolymer length indicates how long the tract of the same nucleotide is after the indel. The colour of a cell indicates how many FN indels of that indel-homopolymer length combination.

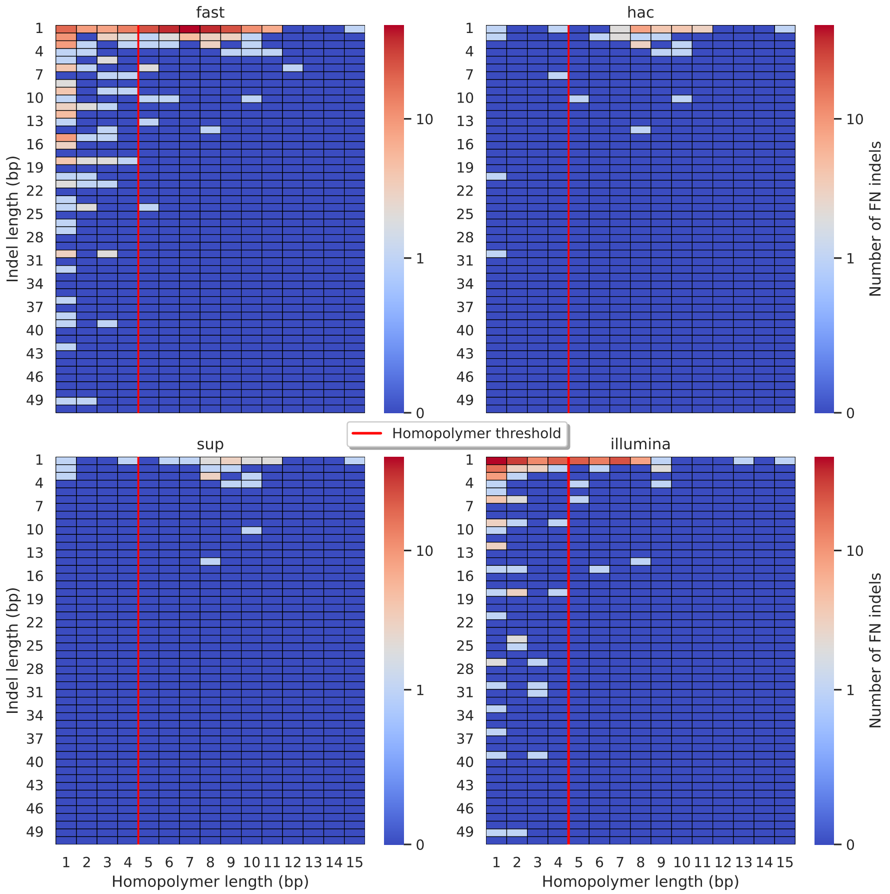


**Figure S19:** Relationship between indel length (x-axis) and homopolymer length (y-axis) for false negative (FN) indel calls for DeepVariant 100x simplex fast (top left), hac (top right), and sup (lower left) calls. Illumina is shown in the lower right for reference. The vertical red line indicates the threshold above which we deem a run of the same nucleotide to be a "true" homopolymer. Indel length is the number of bases inserted/deleted for an indel, whereas the homopolymer length indicates how long the tract of the same nucleotide is after the indel. The colour of a cell indicates how many FN indels of that indel-homopolymer length combination.

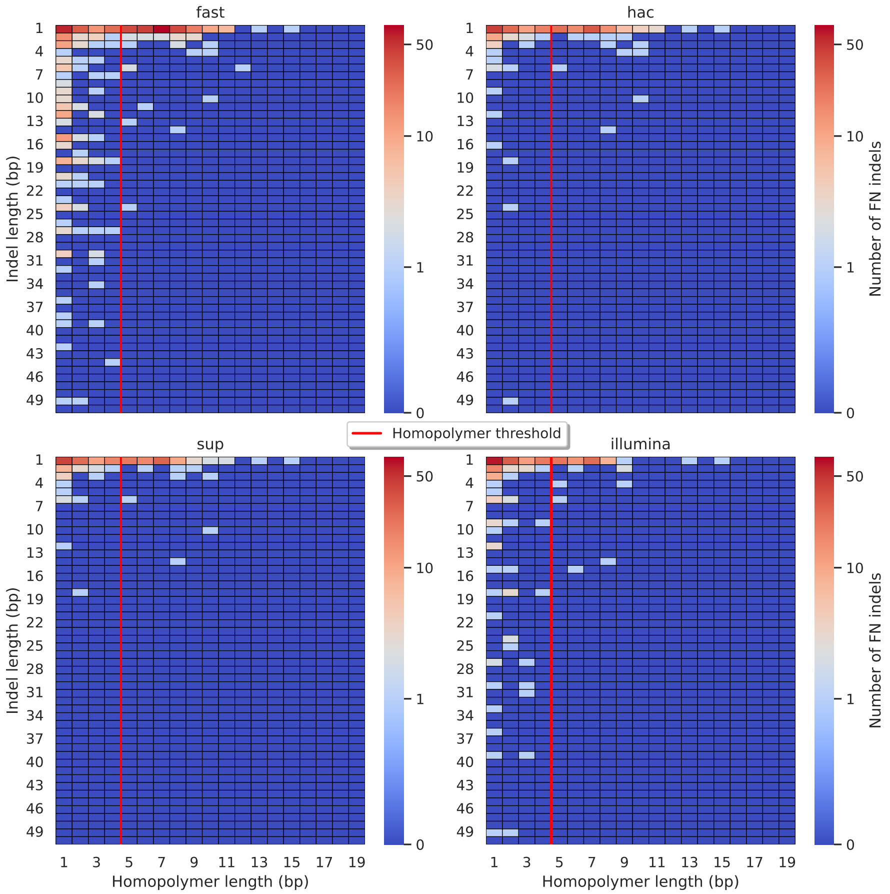


**Figure S20:** Relationship between indel length (x-axis) and homopolymer length (y-axis) for false negative (FN) indel calls for FreeBayes 100x simplex fast (top left), hac (top right), and sup (lower left) calls. Illumina is shown in the lower right for reference. The vertical red line indicates the threshold above which we deem a run of the same nucleotide to be a "true" homopolymer. Indel length is the number of bases inserted/deleted for an indel, whereas the homopolymer length indicates how long the tract of the same nucleotide is after the indel. The colour of a cell indicates how many FN indels of that indel-homopolymer length combination.

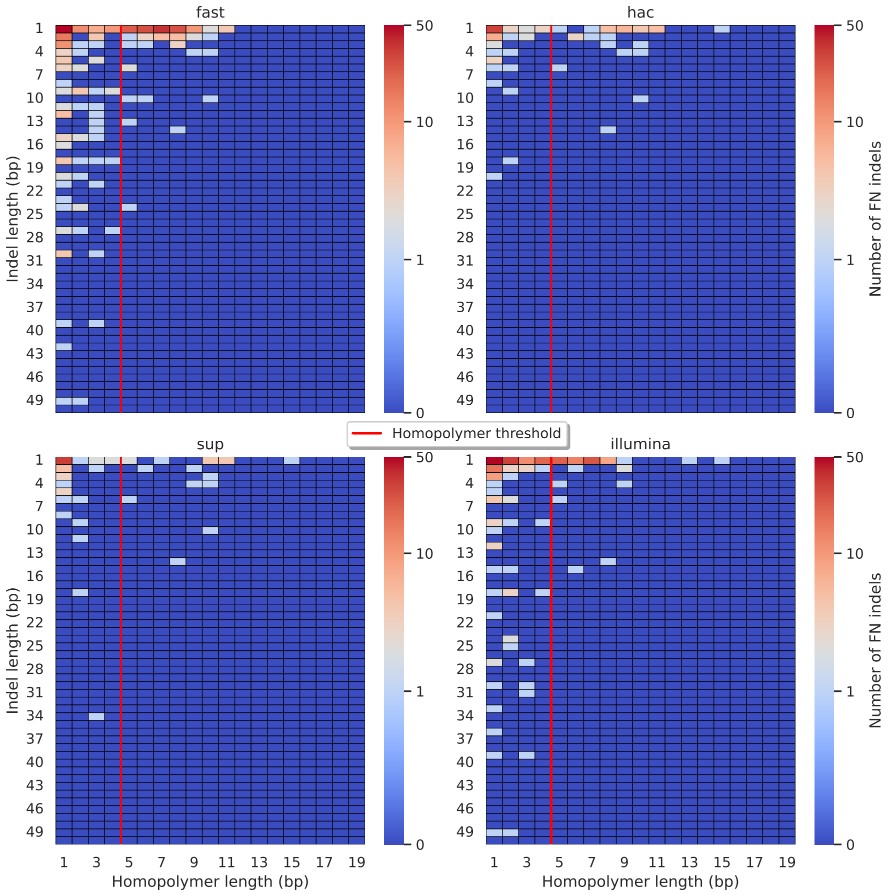


**Figure S21:** Relationship between indel length (x-axis) and homopolymer length (y-axis) for false negative (FN) indel calls for Medaka 100x simplex fast (top left), hac (top right), and sup (lower left) calls. Illumina is shown in the lower right for reference. The vertical red line indicates the threshold above which we deem a run of the same nucleotide to be a "true" homopolymer. Indel length is the number of bases inserted/deleted for an indel, whereas the homopolymer length indicates how long the tract of the same nucleotide is after the indel. The colour of a cell indicates how many FN indels of that indel-homopolymer length combination.

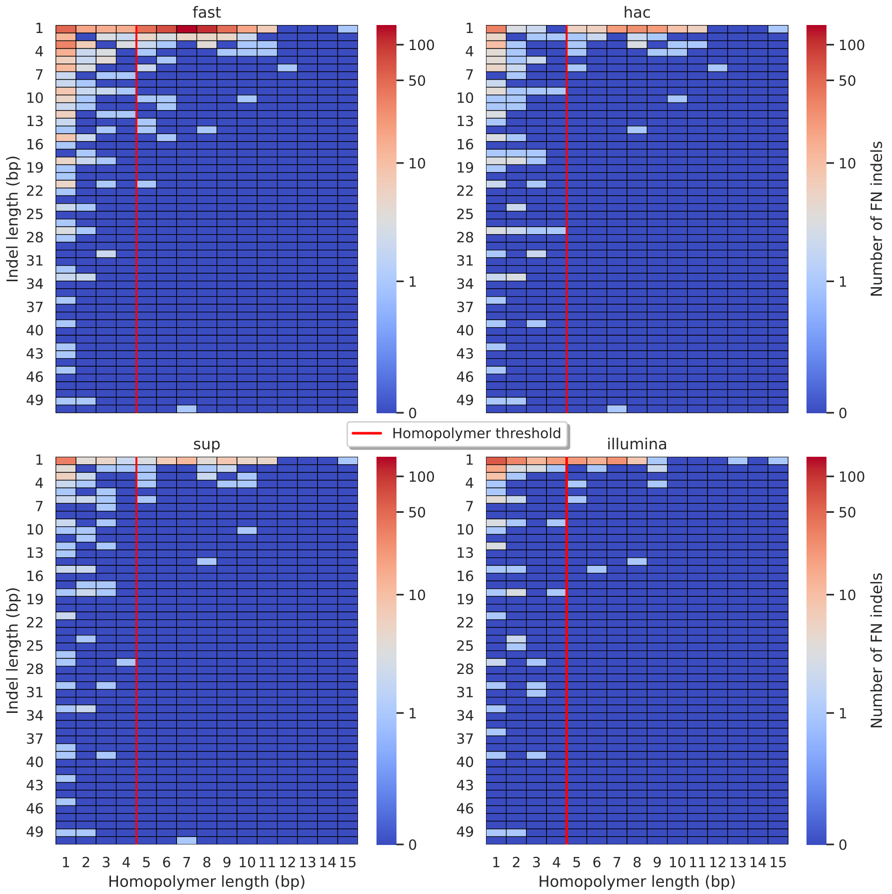


**Figure S22:** Relationship between indel length (x-axis) and homopolymer length (y-axis) for false negative (FN) indel calls for NanoCaller 100x simplex fast (top left), hac (top right), and sup (lower left) calls. Illumina is shown in the lower right for reference. The vertical red line indicates the threshold above which we deem a run of the same nucleotide to be a "true" homopolymer. Indel length is the number of bases inserted/deleted for an indel, whereas the homopolymer length indicates how long the tract of the same nucleotide is after the indel. The colour of a cell indicates how many FN indels of that indel-homopolymer length combination.

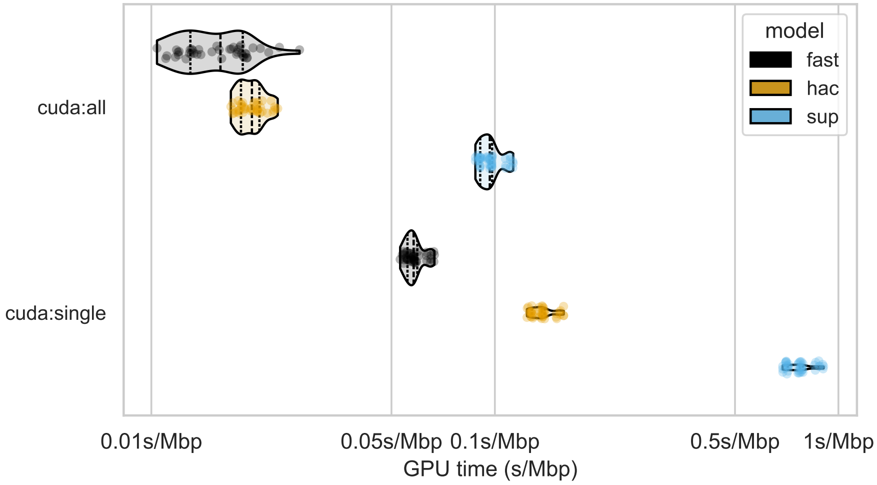


**Figure S23:** Runtime of basecalling ONT data with different models on GPUs. Runtime is presented as seconds per megabasepairs, keeping consistent with Figure 8 from the main text. The y-axis represents basecalling with either 8 Nvidia A100 GPUs (cuda:all) or 1 A100 (cuda:single), with each stratified by basecalling model (colours). Points represent running a single sample-model combination, with each combination being run three times.

### References

[1] James T Robinson et al. "Integrative genomics viewer". In: *Nature Biotechnology* 29.1 (Jan. 2011), pp. 24–26. ISSN: 1546-1696. DOI: 10.1038/nbt.1754.
